# Pleistocene stickleback genomes reveal the constraints on parallel evolution

**DOI:** 10.1101/2020.08.12.248427

**Authors:** Melanie Kirch, Anders Romundset, M. Thomas P. Gilbert, Felicity C. Jones, Andrew D. Foote

**Affiliations:** Friedrich Miescher Laboratory of the Max Planck Society, Max-Planck-Ring 9, 72076 Tübingen, Germany; Geological Survey of Norway, Trondheim, Norway; Evolutionary Genomics, Natural History Museum of Denmark, University of Copenhagen, Copenhagen, Denmark; Department of Natural History, Norwegian University of Science and Technology (NTNU), University Museum, 7491 Trondheim, Norway; Molecular Ecology and Fisheries Genetics Laboratory, School of Biological Sciences, Bangor University, Bangor, UK

## Abstract

Parallel evolution is typically studied by comparing modern populations from contrasting environments, therefore the chronology of adaptive changes remains poorly understood. We applied a paleogenomics approach to investigate this temporal component of adaptation by sequencing the genomes of 11-13,000-year-old stickleback recovered from the transitionary layer between marine and freshwater sediments of two Norwegian isolation lakes, and comparing them with 30 modern stickleback genomes from the same lakes and adjacent marine fjord. The ancient stickleback shared genome-wide ancestry with the modern fjord population, whereas modern lake populations have lost substantial ancestral variation following founder effects. We found modern lake stickleback had lost freshwater-adaptive alleles found in the ancient stickleback genomes, and showed incomplete adaptation, revealing the hitherto underappreciated stochastic nature of selection on standing variation present in founder populations.

**One Sentence Summary:** ‘Pleistocene threespine stickleback genomes reveal insights into the earliest stages of freshwater adaptation’

---

Parallel evolution can occur when natural selection repeatedly acts upon adaptive standing genetic variation, resulting in independently replicated evolutionary outcomes in response to an environmental change (*1,2*). However, a longstanding question in evolutionary biology is whether such parallel evolution repeatedly follows the same genetic pathways to reach convergent phenotypic outcomes, or whether there are constraints preventing parallel processes (*3,4*). Paleogenomic approaches hold the potential to provide new insights into this temporal component of adaptation (*5*), particularly when applied to an emblematic system for the study of parallel evolution such as divergent marine-freshwater pairs of threespine stickleback *Gasterosteus aculeatus* (*6,7,8*).

The threespine stickleback genome is well characterised through mapping, sequencing and transgenics studies, which have provided a strong basis for understanding the role of genes underlying phenotypic changes in skeletal armour, body shape and other morphological and physiological changes associated with freshwater or marine adaptation (*6,9*). Studies of parallel evolution in stickleback typically focus on patterns of genetic and phenotypic variation among contemporary populations and can only track the underlying processes throughout a few decades. Adaptation of marine sticklebacks to freshwater habitats can occur rapidly over tens of generations (*10,11*), facilitated by the reuse of pre-existing genetic variation carried in ancient haplotype blocks within the marine population (*8,12,13*). This standing genetic variation is thought to have been maintained over geological timescales (*12,14*) through recurrent migration between freshwater and marine populations (*2*), and thus repeatedly acted upon by natural selection in freshwater populations. The present adaptive radiation commenced during the transition between the Pleistocene and Holocene epochs, and from glacial to inter-glacial (*6*). Investigating signatures of freshwater adaptation in the genomes of stickleback from this key time point in their evolutionary history has not been possible until now.

Here, we demonstrate the power of paleogenomics to provide novel temporal insights into evolutionary processes by sequencing partial genomes of two 11-13,000-year-old stickleback, which lived during the period immediately after the latest retreat of the Pleistocene Scandinavian Ice Sheet, when many freshwater coastal lakes were forming due to strong post-glacial land uplift. Thus, the ancient samples represent the earliest stages of parallel adaptation to freshwater lakes from a marine-adapted ancestor, thereby providing invaluable insight into the type, availability and maintenance of genetic variation at a formative time in a species adaptive radiation.

## RESULTS

### The ecological and geological context of Late Pleistocene stickleback bones

When the ice sheets retreated after the Last Glacial Maximum, newly formed coastal habitats could be colonised by marine species. Simultaneously, glacio-isostatic rebound, the process whereby the land rises relative to sea-level after being released from compression by ice sheets, resulted in marine basins in many coastal regions being elevated above sea-level (*15*). Once isolated from the sea, these basins gradually changed to a freshwater ecosystem. This ecological change resulted in a distinct sedimentary boundary between a marine sediment facies and a freshwater lacustrine sediment facies which is used by geologists to reconstruct the relative sea-level history (*15*). We examined sediment cores collected from isolation lakes in Finnmark, northernmost Norway (Fig. 1A, fig. S1), an area where post-glacial uplift has caused a net relative sea-level fall of 50-100 m since deglaciation (*16*).

**Fig. 1.**
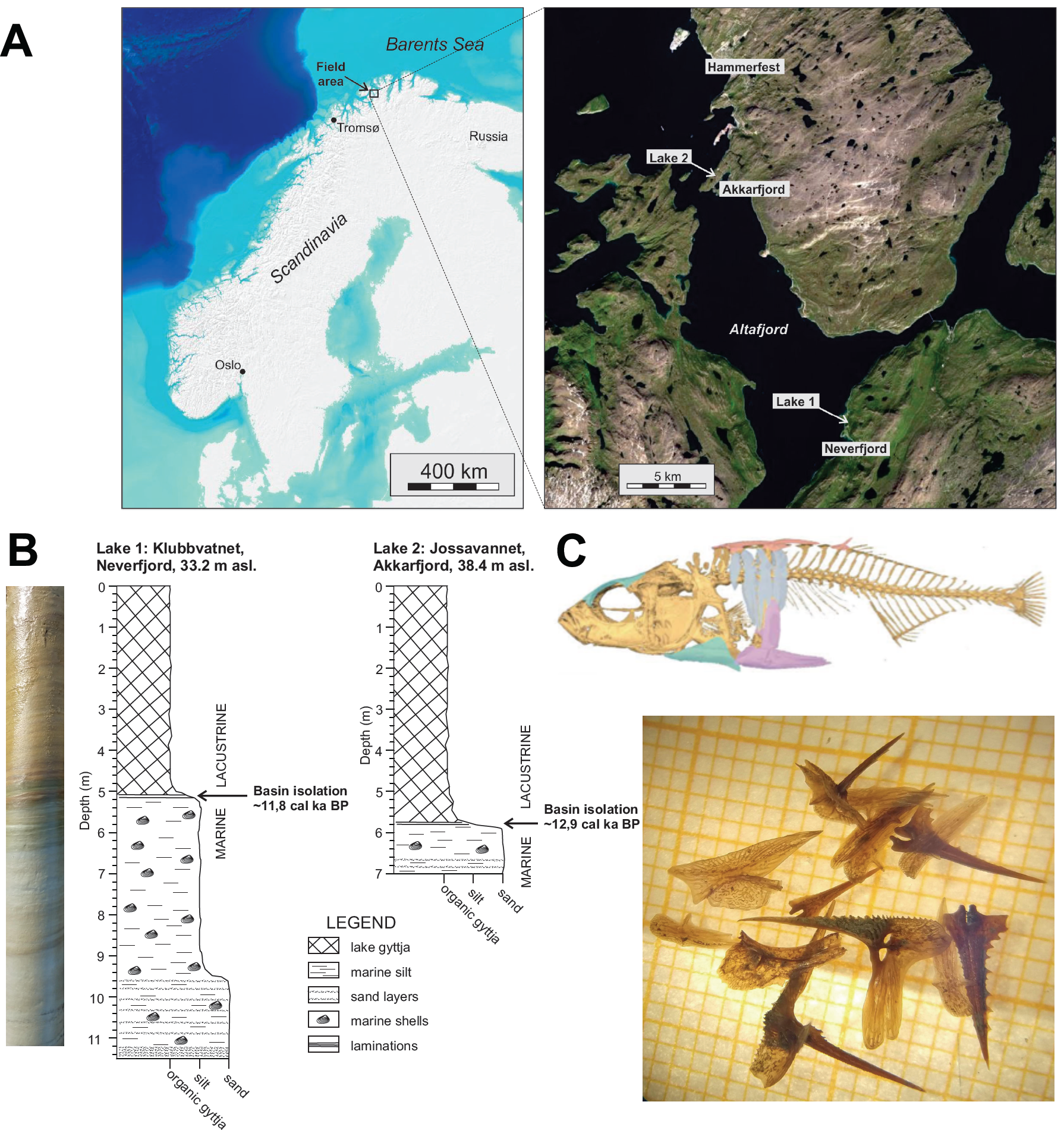
The ecological and geological context of Late Pleistocene stickleback remains. (**A**) Ancient and present-day samples were collected from Klubbvatnet freshwater lake (70° 36’ N, 23° 37’ E; hereafter Lake 1), and from Jossavannet freshwater lake (70° 27’ N, 23° 47’ E; hereafter Lake 2), additionally samples of the marine ecotype were collected from the outer branch area (next to the lake sites) of Altafjord (70° 27’ N, 23° 46’ E). (**B**) The ancient samples were found in the sediment layers of cores from the two lakes corresponding to the isolation phase, dated to ∼11.8 and 12.9 KY BP respectively. An example of the variation in core stratigraphy is shown on the left, schematic diagrams of the stratigraphy in the two study lakes are shown to the right. (**C**) Bones, spines and bony armour plates found in Lake 2. The background grid is mm-scale. Bone positions illustrated on an X-ray scan of modern freshwater fish (above). See fig. S1 for details.

Stickleback bones, spines and bony armour plates were found in the layers of cores from two lakes corresponding to the brackish phase when the lakes became isolated from the marine fjord and were transitioning to freshwater. Radiocarbon dating of organic matter from the same stratigraphic depth placed the age of the stickleback bones as 12,040-11,410 cal yr BP for Lake 1 and 13,070–12,800 cal yr BP for Lake 2 (Fig. 1B). Shotgun sequencing was performed on a stickleback spine from Lake 1, and a bony armour plate from Lake 2 (Fig. 1C, fig. S1). Nucleotide misincorporations relative to the reference genome indicate an approximately 30% deamination rate of cytosine at the read-ends (fig. S2). Such post-mortem damage patterns are characteristic of the degradation in ancient DNA samples that are thousands of years old (*17*). To the best of our knowledge, these are the oldest fish bones from which genomic data have been obtained (*18,19*).

### Ancient lake stickleback share ancestry with modern marine stickleback

To understand the importance of the ancient samples in the chronology of freshwater adaptation, we first established their relationships to present-day sticklebacks sampled from both the local geographic area and more geographically distant locations across the Northern Hemisphere (*8*). Comparison of genome-wide covariance in allele frequencies using Principal Component Analysis (PCA), excluding marine-freshwater parallel divergent regions identified by Jones et al. (*8*), separated out samples by geography into Pacific and Atlantic clusters on PC1 (*P* < 0.001; Fig. 2A, fig. S3). Both ancient samples show closest affinity to present-day Atlantic populations, as expected. The same analysis using only marine-freshwater parallel divergent genomic regions, revealed both ancient samples cluster with the globally sampled marine individuals on PC1 (*P* < 0.005; Fig 2B, fig. S3), suggesting under a polygenic model that the two fish would have had a predominantly marine phenotype.

**Fig. 2.**
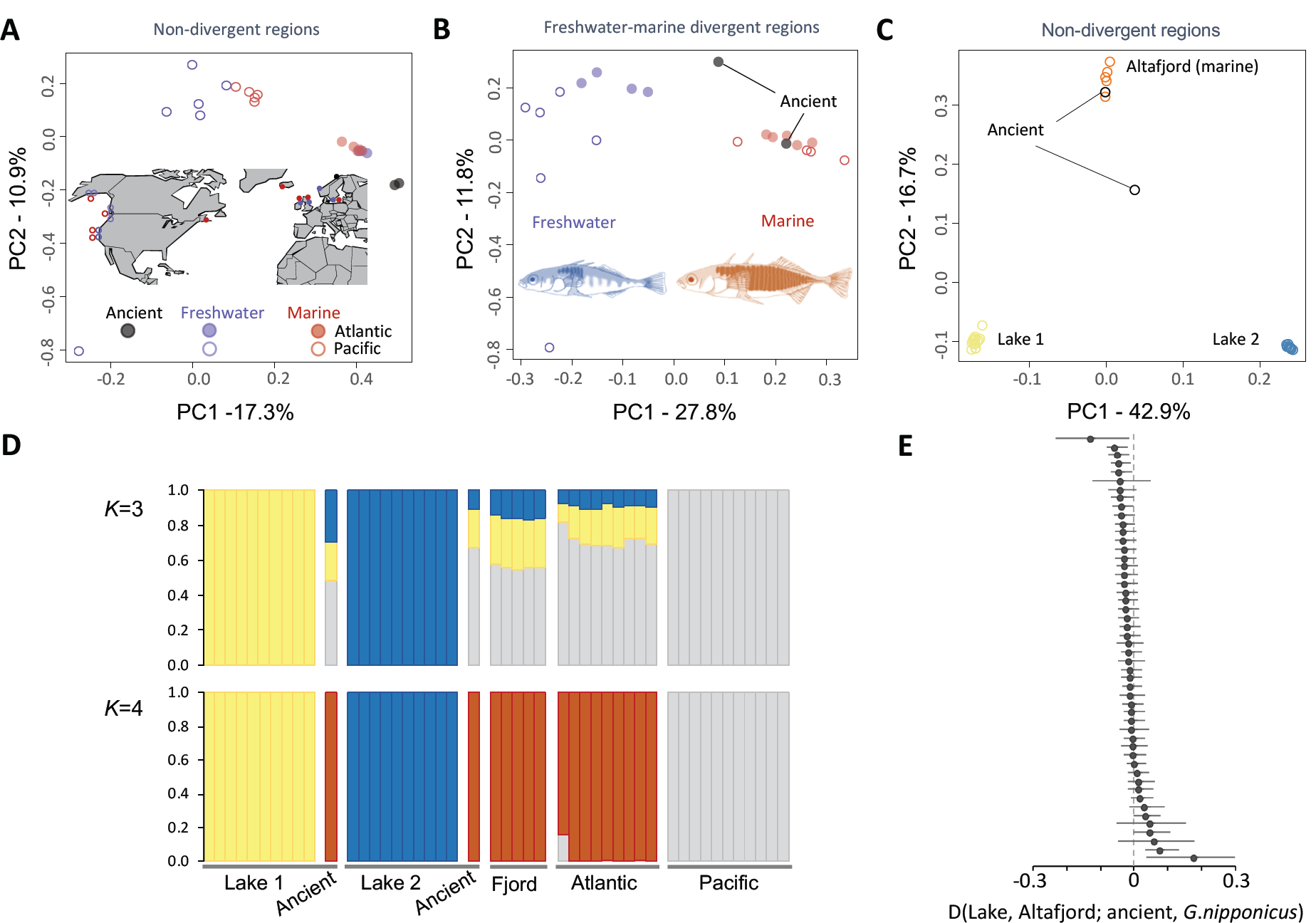
Relationships between ancient and present-day stickleback. Principal component analyses (PCA) of the global dataset from Jones et al. (*8*) and ancient samples based on (**A**) transversions in non-divergent regions, (**B**) transversions in freshwater-marine divergent regions identified by Jones et al. (*8*). (**C**) PCA of local present-day and ancient samples using transversions in non-divergent regions. (**D**) Admixture plots of combined global and local populations. Ancient samples are denoted by asterisks. (**E**) *D*-statistics of the form (Lake 2, Altafjord; ancient, Japan Sea stickleback) testing whether the ancient sample shares more alleles with the present-day lake or fjord samples. The results were not significantly different from zero, suggesting the ancient sample is symmetrically related to both. Error bars show 1 SE.

Focusing on the two focal freshwater lakes from which the ancient samples were recovered, and the adjacent marine fjord, we found strong covariance among present-day genomes within each lake, and likewise among genomes from the fjord (Fig. 2C). The focal lakes in this study are found less than 300 metres from the fjord on land that rises steeply to the post-uplift height of 33.2 and 38.4 metres above sea-level, making subsequent immigration of marine stickleback unlikely. Thus, present-day stickleback in these isolation lakes are likely to be descendants of early colonists that included the ancient samples. Accordingly, the strongest differentiation was between genomes from Lake 1 and those from Lake 2 (PC1, *P* < 0.005), explaining 42.9% of the variance in the data (Fig. 2C; fig. S3). The ancient genomes cluster more closely with the marine fjord samples, though the Lake 1 ancient sample is found between the fjord and lake samples along PC2 (*P* < 0.001; Fig. 2C; fig. S3). A similar pattern is seen when considering the marine-freshwater divergent regions (fig. S4). Clustering patterns in the PCA were reflected in admixture plots, in which both ancient samples shared ancestry components with the present-day fjord samples, whilst present-day lake samples retained just a lake-specific subset of this ancestral variation (Fig. 2D).

The placement of the ancient samples relative to the present-day fjord and lake populations in the PCA, and the pattern of ancestry components in the admixture plots could reflect the strong independent drift in the two lake populations since the death of the two ancient sticklebacks. Estimation of covariance in allele frequencies in the form of an admixture graph, supports this inference and suggests that the ancient samples are approximately equivalently ancestral to the fjord and two modern lake populations, the latter having undergone the greatest drift in allele frequencies (fig. S5). To formally test this hypothesis, we computed *D-*statistics corresponding to the population history *D*(modern fjord, modern lake; ancient lake, *G. nipponicus*). The *D*-statistic tests are consistent with the ancient sample from Lake 2 being symmetrically related to the present-day marine fjord and freshwater Lake 2 populations (−3 < |Z| < 3; Fig. 2E). Thus, the ancient stickleback from Lake 2 is inferred to have lived close to the time of the divergence of the ancestral fjord and lake populations. The *D*-statistics also confirm that strong independent drift in allele frequencies in the two lake populations since their colonisation underlie the patterns observed in the PCA and admixture plots (Figs. 2C & D).

### Founder effects reduce the efficacy of selection on freshwater alleles

Our findings of relative isolation and strong independent drift in the lake populations suggest reduced effective population size (*N*_e_). This has implications for the ability of the population to adapt to freshwater, as the effectiveness with which natural selection fixes advantageous alleles in a population depends not only upon the selection coefficient (*s*) of an allele, but also on *N*_e_ (*20*). To better understand how *N*_e_, and by proxy, the strength of natural selection had varied through time we reconstructed the demographic history of fjord and lake populations using the pairwise sequentially Markovian coalescent (PSMC) method (*21*). We inferred overlapping estimates of *N*_e_ for the fjord and lake populations between 100–20 KY BP. However, demographic histories start to diverge between 20-10 KY BP (Fig. 3A, fig. S6), approximately the time of the directly dated isolation of the two lakes (Fig. 1B). Following isolation from the marine population, we observed a steep decline in inferred *N*_e_ in both lake populations, consistent with that reported by Liu et al. (*22*) for other lake populations.

**Fig. 3.**
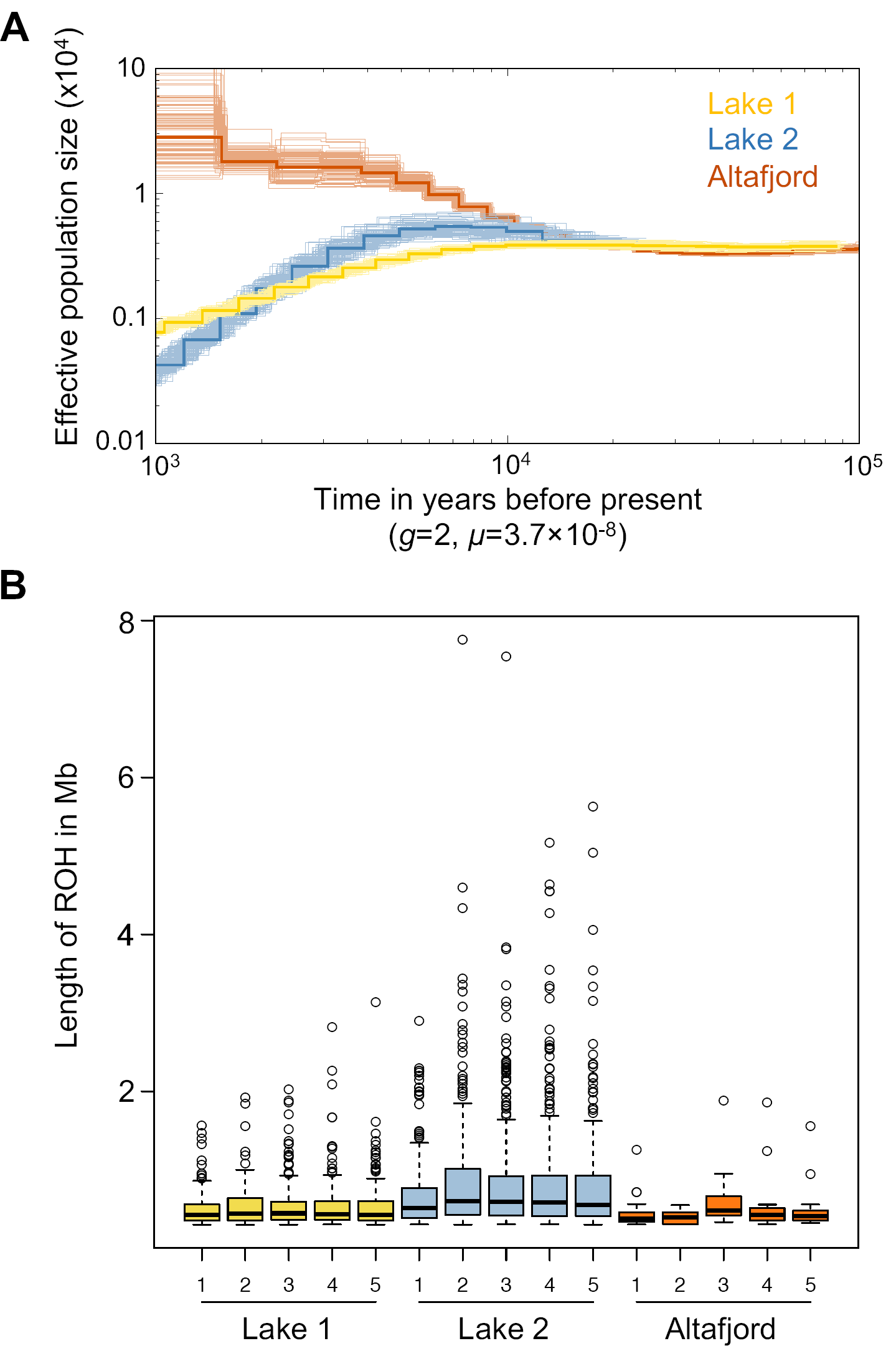
Demographic history of local marine and freshwater stickleback. (**A**) PSMC estimates of changes in effective population size (*N*_e_) over time inferred from the autosomes of a Lake 1 (yellow), Lake 2 (blue) and Altafjord (orange) sample. Thick lines represent the median and thin light lines of the same colour correspond to 100 rounds of bootstrapping. (**B**) Distribution of the length of runs of homozygosity (ROH) greater than 0.3 Mb in the genomes of five samples each from Lake 1 (yellow), Lake 2 (blue) and Altafjord (orange). The thick black line shows the median. The bottom and top of the box represent the 1^st^ (Q1) and 3^rd^ (Q3) quartile. The upper whisker corresponds to the smaller value of the maximum length of ROH or the sum of Q3 and 1.5 times the size of the box (Q3-Q1). All values above the upper whisker are shown as black circles. The lower whisker shows the smallest length of ROH for the corresponding individual.

Runs of homozygosity (ROH) provide further inference of recent demographic history (*23*). The impact of smaller population size of the lake stickleback results in an increased proportion of the genome being identical by descent and in long ROH, particularly in Lake 2 (Fig. 3B). The sum of ROH longer than 300 kb was significantly higher in both lake populations than in the fjord population (Wilcoxon rank sum test, *P* < 0.01). ROH sum up to 83 Mb for stickleback from Lake 1 and 277 Mb for Lake 2 (fig. S7), corresponding to 18% and 60% of the genome respectively, indicating that both populations had undergone genetic bottlenecks. Taken together, our population genetic results indicate strong founder effects and ongoing reduction in effective population size influence the efficacy of selection on freshwater alleles both during and after the colonisation event.

### The accumulation of freshwater adaptive alleles in the ancient and modern samples

In addition to the constraints imposed by reduced effective population size, the progression of parallel evolution will also be dependent upon the availability of freshwater adaptive alleles upon which selection acts. Our paleogenomic data provide the first opportunity to directly compare standing genetic variation present in a freshwater lake at the start of the freshwater adaptation process to present-day genetic variation. Using genomic positions with data present in the ancient stickleback samples, and after down-sampling present-day Lake 2 and Altafjord genomes to equivalent levels, we identified 814 regions of the genome that contain sites with strong divergence among present-day freshwater Lake 2 and marine Altafjord fish (locally divergent regions). At these locally divergent regions, the ancient stickleback from Lake 2 predominantly shared alleles with the present-day fjord population (Fig. 4A). Fixed alleles in these locally divergent regions could represent instances of fixation due to drift in the lake population, rather than having a functional role in freshwater-marine adaption.

**Fig. 4.**
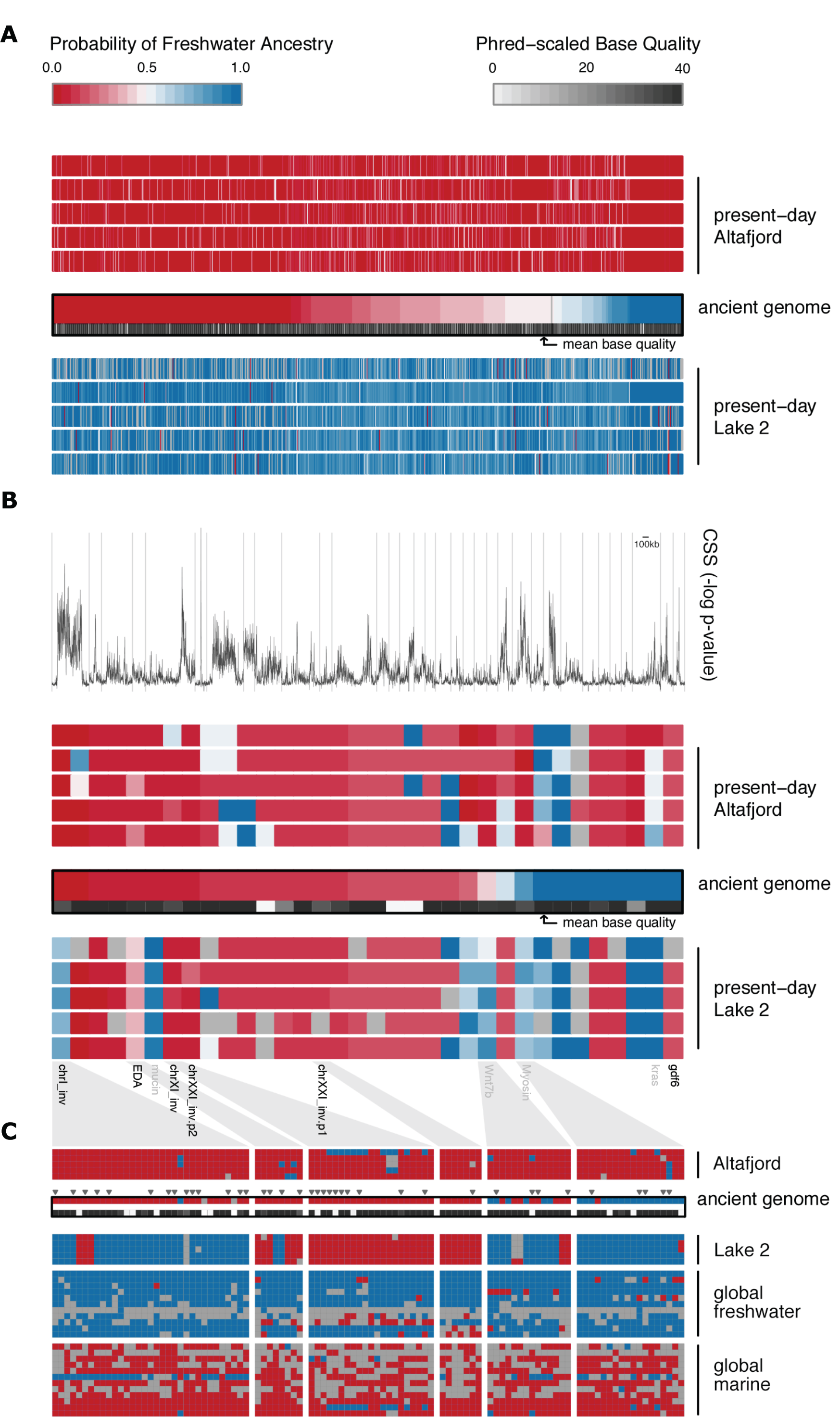
Marine versus freshwater adaptive alleles in the ancient and modern samples. (**A**) The probability of ‘Lake 2 ancestry’ in the ancient sample is plotted for 814 locally-divergent regions of the genome (windows containing variant(s) with fixed allele frequency difference among 5 fish from each of Altafjord and Lake 2). Windows are plotted from left to right according to the probability of freshwater ancestry in the ancient genome. For each genomic region, the mapDamage-rescaled base qualities are plotted in grey-scale for the ancient genome. The probability of freshwater ancestry in five fish from each of present-day Altafjord and Lake 2 populations respectively are shown above and below the ancient genome probabilities. (**B**) The probability of freshwater ancestry in the ancient genome is plotted for 34 genomic regions underlying marine versus freshwater adaptation. These regions were identified based on parallel divergence among global marine versus freshwater populations using cluster separation score (CSS) (*8*). Regions of the genome are plotted from left to right according to the probability of freshwater ancestry in the ancient genome. For each genomic region, the mean mapDamage-rescaled base qualities are plotted in grey-scale below the corresponding probability scores for the ancient genome. The probability of freshwater ancestry in the present day Altafjord and Lake 2 populations are respectively shown above and below the ancient genome probabilities. (**C**) Underlying genotypes at a focal subset of adaptive loci. Rows represent individual fish; columns represent individual single nucleotide polymorphisms; red boxes indicate marine alleles; blue boxes indicate freshwater alleles; grey boxes are missing data. The mean mapDamage-rescaled base qualities are plotted in grey-scale below the corresponding ancient genome probabilities for each site. Transversions, which are less prone to DNA damage due to deamination, are marked by grey triangles.

At genomic regions underlying marine versus freshwater adaptation based on the parallel divergence among a global dataset of marine and freshwater populations (*8*), the ancient genome also carries predominantly marine adapted genotypes, yet carries freshwater genotypes at a greater number of adaptive loci (∼24%) than in the comparison between the local populations (Fig. 4B). The present-day Lake 2 fish carry similar proportions of globally shared marine and freshwater genotypes to the ancient genome suggesting incomplete freshwater adaptation. However, the composition of the adaptive alleles carried by the present-day freshwater stickleback in Lake 2, differ from those found in the ancient genome (Fig. 4B). For example, the ancient genome carries marine versions of the chromosome I inversion harbouring Na+/K+ ion transporter ATPase1a2 and the chromosome II mucin region, whereas the present-day Lake 2 stickleback carry freshwater alleles (Fig. 4C). In contrast, the ancient genome carries freshwater adaptive alleles at some loci where both present-day lake and fjord populations carry marine alleles, e.g. the *GDF6* region, a major effect locus driving bony armour plate size (*24*), on chromosome XX (Fig. 4B), 17.3Mb of chromosome IX and 9.3Mb of chromosome VIII. Therefore, it appears that some freshwater adaptive haplotypes available as standing genetic variation during the founding of Lake 2 have subsequently been lost during the past 12,000 years.

The ectodysplasin (*EDA*) signalling pathway has a key role in the parallel evolution of the low-plated freshwater phenotype due to repeated selection of alleles derived from an ancestral low-plated haplotype (*12,25,26*). There is strong evidence that these alleles persist as standing genetic variation in the marine population (*2,8,12,27*). Consistent with the importance of *EDA* in marine-freshwater phenotypic divergence, we find differentiation between the marine fjord and lake populations (figs. S8, S9, S11), and evidence that both lake populations share the core freshwater haplotype at the EDA locus (fig S11). ROH are prevalent at this locus in the two lake populations (figs. S10, S11), consistent with a ‘hard sweep’ of an extended haplotype under selection in too short a time frame for recombination to restore genetic variation (*28*). Inspecting the underlying genotypes we find extended freshwater haplotypes shared among individuals within each lake, but differing between individuals in Lake 1 and Lake 2 (extending to the right and left flanks respectively, fig. S11). In both populations the ancient freshwater haplotype is flanked by marine haplotypes (fig. S11). This further highlights the independence and associated stochasticity of freshwater adaptation by stickleback in lakes just 25km apart colonised from a shared ancestral genepool.

While studies have reported adaptation to freshwater over decadal timescales by stickleback (*10,11*), by comparing the genotypes of Late Pleistocene stickleback remains to present day genomes we find evidence for a slower and more stochastic process. Our empirical findings are supported by forward simulations (figs. S12-S22), in which freshwater alleles present as low frequency standing variation slowly rise to high frequency after the colonisation of isolation lakes, with limited parallelism between lakes. Our simulations highlight that parallel evolution is constrained by the frequency of freshwater alleles in the founding population, low migration (*29*), and are consistent with our coalescent estimates of changes in effective population size indicating the stochastic loss of freshwater alleles through increased drift during prolonged founder-associated population bottlenecks. The stochastic loss of freshwater-adapted alleles during colonisation of the Atlantic from the Pacific, has been proposed to have reduced parallelism in freshwater adaptation in Atlantic stickleback (*30*). Our results suggest these demographic processes also occur during the colonisation of individual lakes, explaining observations of variation in parallelism of freshwater adaptation globally (*30,31*) and among geographically proximate lakes (*32*). Thus, while the adaptation of threespine stickleback to freshwater is broadly viewed as a highly deterministic process, we find an underappreciated role for stochasticity in this key model system for the study of parallel evolution.

## ACKNOWLEDGEMENTS

We thank Mark Ravinet, Per-Arne Amundsen and Ian Mayer for help with permitting advice. Christian Carøe for advice on extraction and library build. Mike Martin for logistical support. Lina Gislefoss and Thomas Lakeman for help with lake coring. Camilla Scharff-Olsen and Marta Ciucani for ancient DNA lab support. Mette Juul Jacobsen and Lasse Vinner of the National High-throughput DNA Sequencing Centre, University of Copenhagen, Denmark for assistance with Illumina sequencing and associated data processing of ancient samples. Enni Harjunmaa for assigning the ancient stickleback bones to their corresponding position in an X-Ray scan of a modern stickleback. Kavita Venkataramani for DNA extraction and library preparation of the modern samples. Heike Budde, Katrin Fritschi and Ilja Bezrukov for assistance with Illumina sequencing and associated data processing. Katrin Töpner for consultation for creating SLiM simulations. M.A.K. is supported by the International Max Planck Research School “From Molecules to Organisms” and the German Research Foundation (DFG). A.D.F. and the ancient DNA labwork and sequencing were supported by the European Union’s Horizon 2020 research and innovation programme under the Marie Skłodowska-Curie grant agreement No. 663830. The geological field work was funded jointly by NGU and the archaeological research project “Stone Age Demographics” at the University of Tromsø.

