## Supplementary materials for "Pleistocene stickleback genomes reveal the constraints on parallel evolution"

### Materials and Methods

#### Ancient sample collection and geological analysis

Sediment core samples from a number of lakes in the Finnmark region were collected in late spring 2018 and 2019, as part of a study on postglacial relative sea-level changes. Both of the lakes presented in this paper, are relatively shallow and were cored with a "Russian-type" peat corer (*33*). The one metre-long, half-cylinder-shaped samples of lake deposits were collected and transported to the laboratory. As part of the sediment analysis, bulk samples of core material were carefully subsampled and wet-sieved at 125-µm for analysis of macroscopic remains of biota. During basin isolation from the sea, fundamental environmental changes lead to a complete replacement of floral and faunal assemblages, which is apparent in sediment core biostratigraphy (*15*). Marine to lacustrine transitions are often visually distinct as laminated facies and can usually be further determined to within a few centimeters, using preserved remains from certain marine, brackish and freshwater organisms. One such organism which has often been found indicative of a lake in the isolation phase is the three-spine stickleback (*16*,*34*,*35*,*36*). When found, stickleback bones were carefully picked out from the wet-sieved residual material, cleaned and dried at low temperature, before being sent for biological analysis.

#### Modern sample collection and DNA extraction

Adult threespine stickleback specimens were collected using minnow traps from the two lakes from which our ancient stickleback samples originated: Klubbvatnet freshwater lake above the village of Neverfjord (70° 36′ N, 23° 37′ E; Lake 1), and Jossavannet freshwater lake (70° 27′ N, 23° 47′ E; Lake 2). Additionally, samples of the marine ecotype were collected from the outer branch area (next to the lake sites) of Altafjord (70° 27′ N, 23° 46′ E). Samples were collected under permit (201300202-62) from Finnmark Fylkeskommune. Upon sampling, stickleback were euthanized and stored in 95% ethanol. Lab work was performed at the Friedrich Miescher Laboratory of the Max Planck Society in Tübingen. Fin clips of the collected sticklebacks were used for standard Proteinase K digestion (New England Biolabs GmbH, Frankfurt am Main, Germany). Each sample was first incubated for 5 hours at 58°C in 400 µl lysis buffer (50mM Tris-HCl pH=8, 0.1M NaCl, 10mM EDTA pH=8, 0.8% SDS, 15 µg proteinase K) and for 30 min at 37°C with additional 2 µg RNAse A. After adding 150 µl 5M potassium acetate, the sample was stored at 4°C overnight and centrifuged at 4000 rpm for 30 min. Extracted DNA was purified from the supernatant via AmpureXP bead purification (Beckman Coulter GmbH, Krefeld, Germany).

Nextera library preparation was performed with assembled Tn5 bound to magnetic beads. 5 ml of Hydrophilic Streptavidin Magnetic Beads (NEB) were transferred into a falcon tube, placed on a magnet and the bead-storage buffer was removed. The beads were washed with 20 ml streptavidin binding buffer (0.6 M NaCl, 10 mM Tris pH=8.0, 0.5 mM EDTA, 0.1% Triton X-100). Afterwards 30 ml streptavidin binding buffer as well as 400 µl assembled Tn5 were added to the magnetic beads. The mixture was inverted and rotated at 10 rpm at RT for 30 min. The falcon tube was again placed on the magnet, the buffer was removed and replaced by 30 ml of Dialysis buffer (50 mM HEPES-KOH pH 7.2, 0.2 M NaCl, 0.2 mM EDTA, 2 mM DTT, 0.2% Triton X-100, 20% glycerol). The suspension was rotated for another 5 min at 10 rpm, the buffer was removed and 20 ml Dialysis buffer were added for storage of Tn5-on-beads.
2 µl extracted DNA was used for tagmentation with 10 µl Tn5-on-beads in 1x TAPS-DMF buffer (10 mM TAPS, 5 mM $MgCl_2$, 10% DMF) for 15 min at 55°C (total volume 20 µl). 50 µl SDS wash buffer (10 mM Tris pH=8, 30 mM NaCl, 0.1% Triton X-100, 0.3% SDS) were added to the tagmented product for SDS stripping and the suspension was incubated at 55°C for further 4 min. To remove SDS, the samples were placed on a magnet, the supernatant was removed and the tagmented DNA bound to the beads was washed twice with wash buffer (10 mM Tris pH=8, 30 mM NaCl, 0.1% Triton X-100).The library was dual indexed and amplified in a 9-cycle PCR using Q5 High Fidelity Polymerase (Biolabs New England). 50 µl PCR-Mastermix (200uM dNTPs, 300uM of each nextera primer, 1 U Q5 High Fidelity Polymerase, 1x Q5 buffer) was therefore added to each sample. PCR temperature profile included an initial step at 72°C for 5 min to fill up the 9 basepair long gaps made by Tn5 during the tagmentation, an activation step at 98°C for 30 s, followed by 9 cycles of denaturation at 98°C for 15 s, annealing at 65°C for 20 s and elongation at 72°C for 90 s. Amplified DNA was then purified with AmpureXP bead purification (Beckman Coulter GmbH, Krefeld, Germany).

**Ancient DNA lab work**

Ancient DNA lab work was conducted at the dedicated ancient DNA facilities at the Centre for GeoGenetics, University of Copenhagen. DNA was extracted using a silica-based method, where each individual bone or spine was incubated overnight under motion at 55°C in 500 µl extraction buffer (0.45 M EDTA, 0.1 M UREA, 100 µg proteinase K). Each sample was then centrifuged at 2300 rpm for 5 min and the supernatant was collected and concentrated and purified using a Zymo-Spin V reservoir (Zymo Research Irvine, CA, USA) and Qiagen MinElute spin column (Qiagen, Inc., Valencia, CA, USA). To maximise library complexity by reducing the number of DNA purification steps during library preparation, an Illumina library was constructed using the blunt-end single tube (B.E.S.T.) method (*37*). The library was dual indexed and amplified in either a 15-cycle (Lake1 sample) or 20-cycle (Lake 2 sample) PCR using AmpliTaq Gold (ThermoFisher Scientific). The 50ul PCR reaction contained 15ul of library, 25uM dNTP, 1x PCR buffer, 2.5 mM $MgCl_2$, and was made up to 50ul with molecular grade water. PCR temperature profile included an activation step at 95°C for 5 min, followed by 15 cycles of denaturation at 95°C for 30 s, annealing at 55°C for 30 s and elongation at 72°C for 1 min, with a final extension step at 72°C for 7 min. PCR products were then purified using Agencourt AMPure XP beads (BeckmanCoulter). The dual index amplified library of the Lake 1 sample had mean insert size of 186bp, including 114bp of index and adapters, indicating mean DNA fragment size of 72bp. Peak molarity was 286 pmol/l, comprising 34% of the library (including adapter dimer, lower and upper size markers) as quantified using an Agilent 2200 TapeStation instrument with D1000 High Sensitivity ScreenTape and reagents. The dual index amplified library of the Lake 2 sample had mean insert size of 186bp, including 114bp of index and adapters, indicating mean DNA fragment size of 72bp. Peak molarity was 26,100 pmol/l, comprising 85% of the library (including lower and upper size marker peaks). Extraction, library build and index PCR blanks were also included to evaluate potential contamination during the library building process. The Lake 1 DNA library was then pooled with an ancient killer whale *Orcinus orca* DNA library and sequenced across two lanes of 80bp-SE sequencing of an Illumina HiSeq4000, and the Lake 2 library was run across an entire single lane of 80bp-SE sequencing of an Illumina HiSeq4000 at the Danish National High-throughput Sequencing Centre of Copenhagen University (seqcenter.ku.dk).

**Mapping, filtering and masking**

Sequencing data from modern samples generated for this study were demultiplexed and adapters removed using bcl2fastq. Sequencing data of the 21 global samples from Jones et al. (*8*) were downloaded from NCBI in sra format (SAMN00627549-SAMN00627550; SAMN00627914-SAMN00630301). Sra files were transformed to fastq files by using fastq-dump. Sequencing of the ancient samples resulted in 1,381,302 reads for Lake 1 and 627,366,889 million reads for Lake 2. AdapterRemoval removed adapters from the single-end reads of the ancient samples of Lake 1 and Lake 2 and trimmed both Ns and low-quality bases from the reads. The generated fastq files of the 21 global samples, as well as trimmed sequencing data of both ancient samples, were aligned against the reference genome gasAcu1 (*8*) by using the Burrows-Wheeler Alignment Tool with the aln algorithm (*38*), disabling seeding (option‘-l 1024) to turn-off seeding thereby increasing mapped data by including reads with post mortem damage at the read ends (*39*). Sequencing data aligned to the stickleback reference genome encompasses 9,127 reads for the ancient sample from Lake 1 and 7,199,676 reads for the ancient sample from Lake 2. The resulting bam files were sorted and merged by samtools (*40*). Sequencing data of all modern samples were subsequently aligned against the reference genome gasAcu1 (*8*) using the bwa mem algorithm (*38*). Generated bam files were then sorted by samtools (*40*). All types of duplicates in all sorted bam files were identified by MarkDuplicates from Picard Tools [http://broadinstitute.github.io/picard/]. Masked regions as well as the sex chromosome (chrXIX) were removed from the bam files. Masked regions encompassed interspersed repeats and low complexity DNA sequences detected by RepeatMasker (*41*) covering 3.72% of the stickleback genome as well as highly repetitive DNA sequences detected by WindowMasker (*42*) from the NCBI C++ toolkit covering 25.59% of the stickleback genome using -sdust true as setting. After removing duplicates and masked regions, 5,054 reads were left for the ancient sample of Lake 1 and 329,226 reads for Lake 2.

**Assessing postmortem DNA damage and contamination**

Analyses of potential nucleotide misincorporations using PMDtools (*43*) to compare with the modern reference genome revealed that sequencing reads exhibited characteristic post-mortem damage patterns (*17*,*44*), specifically an excess of C→T transitions at the 5′ termini as expected from deamination, and the complementary G→A transitions at the 3′ termini. Therefore, except where otherwise stated, only transversions were considered in downstream analyses that included the ancient samples. Contamination from present-day DNA can be estimated from the number of heterozygous calls in haploid markers (*45*). As both our ancient stickleback were females (see below), contamination could only be inferred from the mitochondrial genomes. Coverage was 1x across most sites, and we detected no heterozygous genotypes in either fish. Furthermore, all segregating sites conformed to the expected haplotype structure based on comparison with modern samples in the same clade. Taken together with the high proportion of reads with DNA damage patterns at the read ends, these results suggest our genome data represented endogenous DNA from the ancient stickleback bones.

**Sexing**

Sticklebacks have an XY sex-determination system, where males are the heterogametic sex. Males are therefore haploid for the X chromosome and diploid for the autosomes, while females are diploid for both the X chromosome and autosomes. The sex of the ancient samples was determined by comparing the mean coverage of the autosomes (excluding unassembled scaffolds), the pseudoautosomal region (chrXIX:1-3300000 and chrXIX:12270000-20240660), and the sex-determining region (chrXIX:3300000-12270000) among 4 female and 4 male modern genomes down-sampled to comparable coverage with each ancient genome. Both ancient samples had approximately equivalent coverage across all three regions implying that both are diploid for the sex-determining region of the X chromosome, i.e. are female (fig. S23). For each modern individual the down-sampling was repeated 10 times with different random seeds to initiate the down-sampling.

**Principal Component Analysis**

We used pseudo-haploid genotype calls of globally distributed modern marine and freshwater genomes (*8*) and the ancient samples. First, we sampled autosomal regions outside of known freshwater-marine divergence associated regions as identified by Jones et al. (*8*) to compare across geographically informative markers. Then, we compared covariance within freshwater-marine divergence associated regions among samples using ecology informative markers. The ancient samples were included in the PC computations and not projected onto PCs of modern samples, which has the advantage of providing a quality control measure. For example, if the ancient samples were impacted by sequencing- or sequence data processing errors, the samples would appear as outliers in the PCA. Instead they cluster with Atlantic and marine samples respectively in the first two PCA plots (Fig. 2A). In the first PCA, both ancient samples were at the extreme edge of PC1 showing closest affinity to the Atlantic samples. Inclusion of a single randomly selected modern sample from Altafjord, Lake 1 and Lake 2 showed that the modern samples also cluster at the edge of PC1 (fig. S24). This suggests structure among the Atlantic samples, potentially reflecting past biogeographical processes (*46*) and gene flow during the Holocene between Pacific and Atlantic lineages, rather than artefacts of unmasked DNA damage or missingness of the data in the ancient samples. Additional filtering steps included in these analyses were the removal of regions of poor mapping quality ($Q < 30$), removal of sites with low base quality scores ($q < 20$), calling only SNPs inferred with a likelihood ratio test (LRT) of $P < 0.000001$, a minimum allele frequency of 0.05 so that alleles had to be called in a minimum of two individuals, discarding reads that did not map uniquely, adjusting q-scores around indels, adjusting mapping quality to 50 for excessive mismatches, discarding bad reads (flag >=256), and the removal of transitions to avoid bias from C→T and A→G DNA damage patterns. The eigenvectors from the covariance matrix were generated with the R function "eigen", and significance was determined with a Tracy-Widom test (*47*) performed in the R-package AssocTest (*48*) to evaluate the statistical significance of each principal component.

The relationship of the 30 modern samples collected from the two lakes and fjord in Finnmark, and the two ancient samples was explored using PCAngsd, a Principal Component Analysis for low depth next-generation sequencing data using genotype likelihoods, thereby accounting for the uncertainty in the called genotypes which is inherently present in low-depth sequencing data (*49*). We restricted the analyses to sites covered in at least one of the two ancient samples, which restricted the dataset to 2,267 SNPs in autosomal chromosomes and outside of freshwater-marine divergent regions. Over 90% of the included SNPs were >100,000 bp apart (fig. S25), thus reducing autocorrelation in covariance due to linkage disequilibrium. Filtering steps were as specified above. The statistical significance of each principal component was estimated from the eigenvalues as above.

**Individual Assignment and Admixture Analyses**

An individual-based assignment test was performed using NGSadmix (*50*), a maximum likelihood method that bases its inference on genotype likelihoods. The input genotype likelihood values were the same as those use above for PCAngsd, as were the filtering steps. NGSadmix was run with the number of ancestral populations $K$ set from 2–10. For each of these $K$ values, NGSadmix was re-run five times for each value of $K$, and with different seeds to ensure convergence. The uppermost hierarchical level of structure, inferred from the greatest step-wise increase in log likelihood, $\Delta K$ (*51*), identified two clusters. PCA and individual assignment-admixture models draw inference from similar information and therefore generate similar axes of variation (*52*,*53*). Both methods typically identify the samples with the greatest population-specific drift that therefore share derived alleles that were rare in the source populations or have lost ancestral alleles from standing variation, as the major axes of structure (*52*). Accordingly, both PCAngsd and NGSadmix identified the uppermost hierarchical level of structure within our data set as being between the Lake 1 and Lake 2 with other samples sharing ancestry with both. However, the $\Delta K$ method is prone to over- or under-estimating population genetic structure (*54*) and performs poorly under scenarios with migration among populations at inferring hierarchical population structure (*55*). Changes in likelihoods show the biggest jumps between $K$=2 and $K$=3, and between $K$=3 and $K$=4, before plateauing to a gradual rising rate in likelihood. Likelihood estimates for $K$=3 and $K$=4 were also highly consistent, having the lowest standard deviations. We therefore present the admixture plots for $K$=3 and $K$=4 in Fig. 2D.

### *D*-statistics

To investigate whether pairs of modern stickleback populations evenly shared derived alleles with the ancient samples, we estimated D-statistics. The test can be used to evaluate if the data are inconsistent with the null hypothesis that the tree (((Lake, Fjord), ancient), outgroup) is correct. The definition used here is from Durand, Patterson, Reich, and Slatkin (*56*):

$$D = \frac{\text{nABBA} - \text{nBABA}}{(\text{nABBA} + \text{nBABA})}$$

where in the tree given above, nABBA is the number of sites where lake and ancient samples share a derived allele, and the fjord samples has the ancestral allele; and nBABA is the number of sites where the fjord and ancient samples share a derived allele, and the lake sample has the ancestral allele. Under the null hypothesis that the given topology is the true topology, we expect an approximately equal proportion of ABBA and BABA sites and thus D = 0. The significance of the deviation from 0 was assessed using Z-scores, which are based on the assumption that the D-statistic (under the null hypothesis) is normally distributed with mean 0 and a standard error achieved using the jackknife procedure. The tests were implemented in ANGSD (*57*) and performed by sampling a single base at each position of the genome to remove bias caused by differences in sequencing depth at any genomic position, removing transitions to avoid bias from C→T and A→G DNA damage patterns, and only considering sites covered in the ancient sample (only the Lake 2 sample was included). Further filtering steps were as specified above for PCA. The Japan Sea stickleback *Gasterosteus nipponicus* (NCBI: PRJDB5176) was used as the outgroup.

### Mitochondrial DNA analysis

Both ancient samples were found to be female based on comparable read coverage across the autosomal, pseudo-autosomal and sex determining regions of the X-chromosome. Therefore, we reconstructed the mitochondrial DNA phylogeny to test whether the ancient samples were directly matrilineally ancestral to either local marine or freshwater populations (fig. S26). The alignment of the mitogenomes of the 81 individuals included in the study by Liu et al. (*22*) was accessed via the dryad repository https://doi.org/10.5061/dryad.46fb1. Modern genomic data from our two lake and fjord study sites were then mapped to the de novo reference mitogenome sequence generated by (*22*), which excluded the control region due to the high number of indels. Mapping criteria were the same as for the nuclear genome. We then assembled a consensus sequence of the mitochondrial genome of each ancient sample, manually inspecting mapped reads and removing those which had a T within 30 bp of the 5′ end where all other sequences had a C at the same site, and similarly removing those which had an A within 30 bp of the 3′ end where all other sequences had a G at the same site. As per Liu et al. (*22*), the mitogenome of *G. wheatlandi* (GenBank Accession no. AB445129) was added as an outgroup and a Maximum Likelihood tree was constructed using PHYML 3.0 (*58*), applying the GTR model for substitution, allowing variable proportions of invariable sites and mutation rates across sites (GTR + I + gamma) and using 100 bootstraps.

The present-day freshwater sticklebacks formed two monophyletic clades corresponding to samples from Lake 1, and those from Lake 2, indicating a recent common maternal ancestor within each lake, and no detectable immigration into either lake (fig. S26). Mitochondrial genomes generated from the present-day marine samples collected from Altafjord, and the ancient samples also fell within the clade representing the so-called 'European Lineage' (*59*), but were intermixed among sequences from Denmark, Germany, Greenland and southern Norway. Thus, the ancient samples did not represent the direct mitochondrial ancestor of the local present-day freshwater population within the same lake. However, mitochondrial DNA represents a single genealogy and the ancient is a single sample, and the high haplotype diversity in our Altafjord sample set, and the lack of fine-scale phylogeographic patterns, suggests the ancestral marine founders of each lake population may have had similarly high mtDNA haplotype diversity.

**Pairwise sequentially Markovian coalescent**

The PSMC model estimates the Time to Most Recent Common Ancestor (TMRCA) of segmental blocks of the genome and uses information from the rates of the coalescent events to infer $N_e$ at a given time, thereby providing a direct estimate of the past demographic changes of a population (*21*). We selected the highest coverage modern genomes, building a consensus sequence of each bam file in fastq format sequentially using: firstly, SAMtools mpileup command with the –C50 option to reduce the effect of reads with excessive mismatches; secondly, bcftools view –c to call variants; lastly, vcfutils.pl vcf2fq to convert the vcf file of called variants to fastq format with further filtering to remove sites with less than a third or more than double the average depth of coverage and Phred quality scores less than 30. Furthermore, we excluded marine-freshwater divergence associated regions and 100 kb flanking either end. The PSMC inference was then carried out using the recommended input parameters as previously applied to threespine stickleback genomes (*22*), i.e. 25 iterations, with maximum TMRCA (Tmax) = 15, number of atomic time intervals (n) = 64 (following the pattern (1*4 + 25*2 + 1*4 + 1*6), and initial theta ratio (*r*) =5. Plots were scaled to real time as per (*22*), assuming a generation time of 2 years and a neutral autosomal mutation rate of $3.7\times10^{-8}$ substitutions/nucleotide/generation. To check for variance in $N_e$ across the genome, we performed 100 bootstrap replicates, conducted by randomly sampling with replacement 5-Mb sequence segments obtained from the consensus genome sequence. To be assured that the inferred demographic history is not impacted by the sequence coverage, sequencing data of freshwater stickleback BS65 from Feulner et al. (*60*) was processed identically to our samples and downsampled to different coverages (fig. S27). The PSMC plot of these samples shows a consistent reduction in $N_e$ for all samples, which suggests that lower coverage can affect the quantitative estimate of $N_e$, but not the overall pattern.

**Runs of homozygosity**

Runs of homozygous genotypes (ROH) were identified using the window-based approach implemented in PLINK v1.07 (*61*) from an input file of genotype likelihoods generated by ANGSD (*57*) with the following filtering settings: removing reads of poor mapping quality (MAPQ < 30), removing sites with low base quality scores (q < 20), calling only SNPs inferred with a likelihood ratio test of P < 0.000001, discarding reads that did not map uniquely, adjusting q-scores around indels, adjusting minimum quality score to 50 for excessive mismatches, and discarding bad reads (flag >=256). We estimated ROH from pruned and unpruned data and found minimal qualitative difference with our data. Sliding window size was set to 300 kb, with a minimum of 50 SNPs at a minimum density of 1 SNP per 50 kb required to call a ROH. To account for genotyping errors, we allowed up to 4 heterozygote sites per 300 kb window within called ROHs, as per ref. (*62*). A length of 1,000 kb between two SNPs was required in order them to be considered in two different ROHs.

**$F_{ST}$ statistics**

$F_{ST}$ statistics between two populations at a time were performed with vcftools (*63*) from an input file in variant call format (VCF) created by bcftools mpileup and bcftools call (*40*)(*64*). The reference stickleback genome gasAcu1 (*8*) and modern samples sequenced at high coverage with five samples from each location were used as input for *bcftools mpileup*. The output file was subsequently piped into *bcftools call* which used the multiallelic-caller to output a VCF file for all fifteen modern samples. Thereafter, the resulting VCF file was used for calculating $F_{ST}$ estimate per site between each two populations based on Weir and Cockersham's method (*65*) by using *vcftools*. The $F_{ST}$ estimate was processed in *R* (*66*) and plotted in 10kb sliding windows with 5kb steps in figures S8-10. For figure S11, the goal was to visualise variation at a finer-scale, we therefore opted to estimate $F_{ST}$ per site. The VCF file was filtered with *vcftools* for sites with a minor allele frequency greater than or equal to 0.07, at least 80% of the data non-missing, a quality value above 30, and mean depth values as well as genotypes with a depth between 2 and 9 excluding indels. Subsequently, $F_{ST}$ was estimated per site (*67*), and plotted in R using Friedman's 'super smoother' with a span of 1/50 (*68*,*69*).

**Comparison of ancient and modern genotypes**

In order to investigate the progression of the ancient genomes towards a freshwater genotype due to natural selection we estimated the probability of sharing local lake versus fjord ancestry, and globally shared marine versus freshwater ancestry in regions associated with geographically local differentiation and with parallel marine-freshwater adaptation respectively. We estimated genotype probabilities only for the Lake 2 ancient sample for which there was sufficient coverage in known freshwater-marine adaptive regions. To maximise the available data we rescaled base quality scores using mapDamage 2.0 (*70*) which penalises the quality score of bases likely to be impacted by post-mortem damage based on the posterior distribution of damage-associated parameters, thereby allowing the inclusion of transitions (with a measure of confidence in the inference drawn from individual bases). First, we focused on regions that were differentiated between the local Lake 2 and Altafjord present-day populations, down-sampled to equivalent coverage to the ancient sample; specifically, selecting windows with at least one fixed difference between the five highest coverage modern samples from each of Lake 2 and Altafjord. The probability of having either 'Lake' or 'Fjord' ancestry is reported for these windows. Next, we investigated the probability of whether sites in the ancient sample and local modern samples had 'marine' or 'freshwater' ancestry in regions previously identified as underlying parallel marine-freshwater adaptation in a global dataset (*8*) Higher probabilities of either marine or freshwater ancestry in individual fish typically reflected the presence of haplotype blocks associated with either habitat, confirmed by visualising at the level of individual genotypes for key adaptive loci.

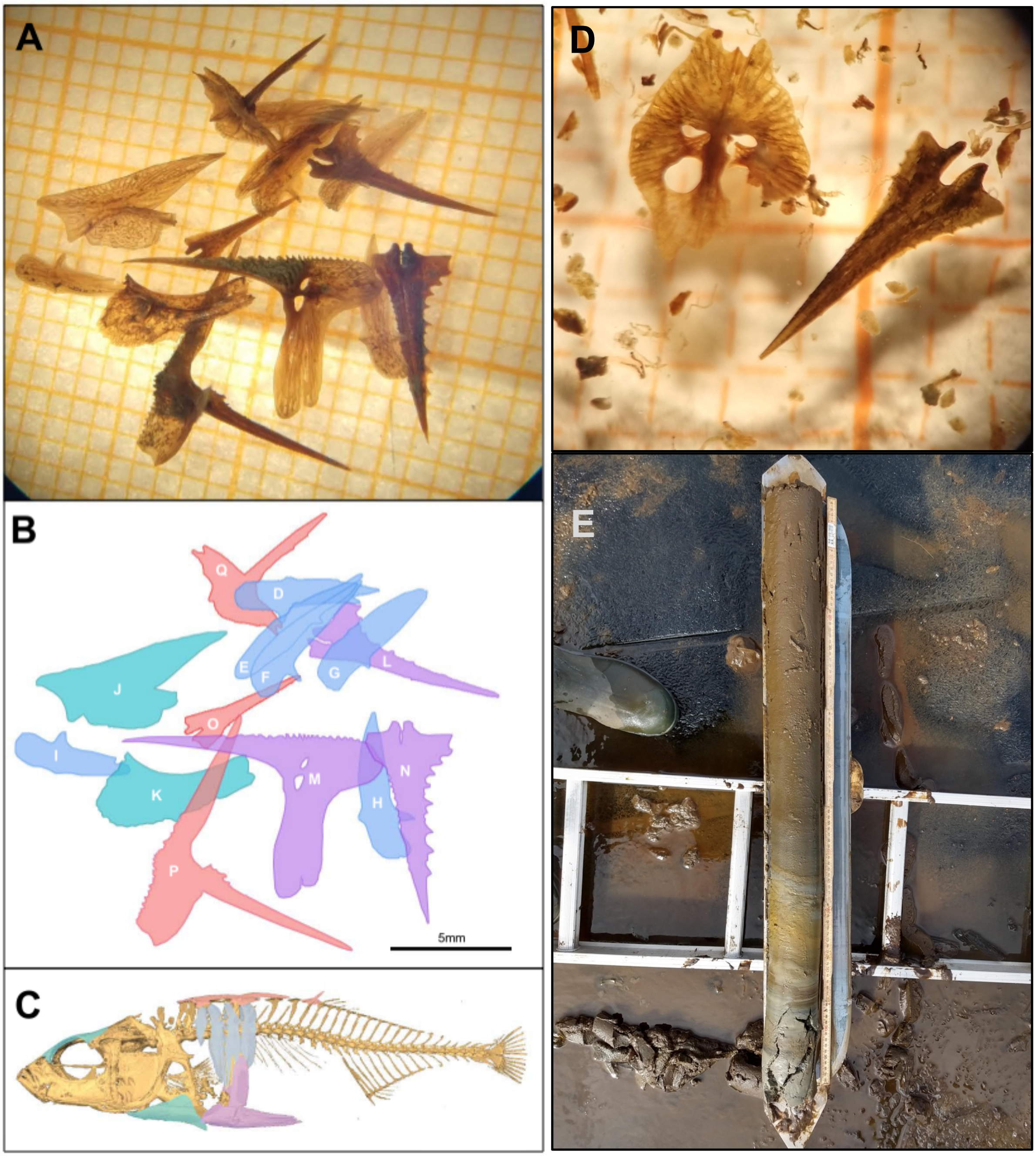


**Figure S1: (A)** Photograph of bones from Lake 1 (Klubbvatnet) near Neverfjord, Finnmark, Norway. Small squares on the backing paper are 2mm$^2$. **(B)** Illustration of bone identities from **(A)**. D-I (blue) are lateral plates. J and K (turquoise) are from the head area. J is the plate-like dermal bone between the gills and the pelvis (ectocoracoid). K is the part of the skull plate that is above the eye (frontal). L, M and N (violet) are the pelvic bones: L and N are the pelvic spines, M is the spine-like element between them and part of the lateral protrusion. O, P and Q (red) are dorsal spines; O has lost its basal plate, while P and Q retain theirs. Scale bar in lower right corner is 5mm. **(C)** Positions of bones from **(A & B)** illustrated on an X-Ray scan of a modern freshwater fish. **(D)** Photograph of bones from Lake 2 (Jossavannet) near Hammerfest, Finnmark, Norway. Small squares on the backing paper are 2mm$^2$. (E) Sediment core from Lake 2. The marine phase of the core is on the right-side of the photograph, which is characterized by marine clay, silt and sand, which transitions to clay gyjttja with laminations during the phase where the lake is partially isolated with occasional marine contact, and then to freshwater gyttja as the lake became fully isolated.

**A**

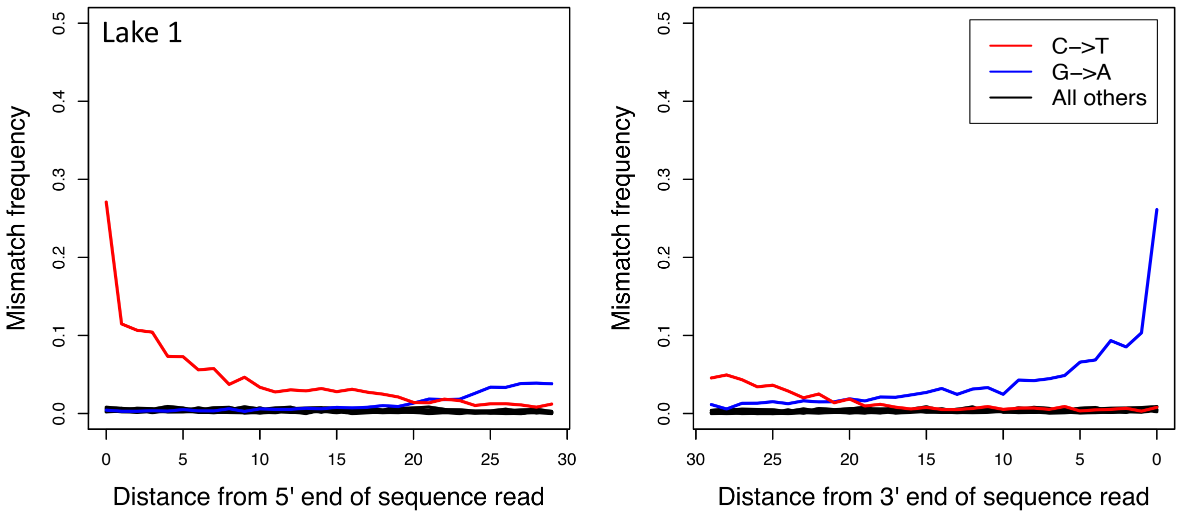


**B**

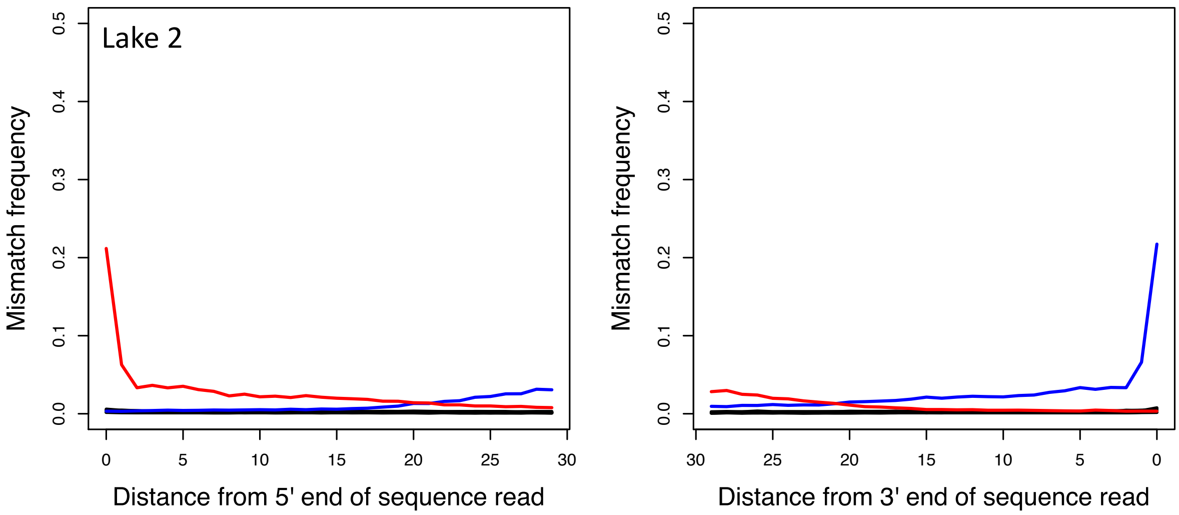


**Figure S2:** Postmortem damage pattern for **(A)** Lake 1 and (**B**) Lake 2. DNA misincorporation errors relative to the 5′ and 3′ read termini sequence data generated from the ancient samples relative to the modern stickleback reference (gasAcu1). The two distributions for post mortem damage signatures (C>T and G>A) are shown in red and blue respectively. All other substitutions are shown in black. Nucleotide frequencies are shown for 30 bases upstream and downstream of the 5′ and 3′ read termini.

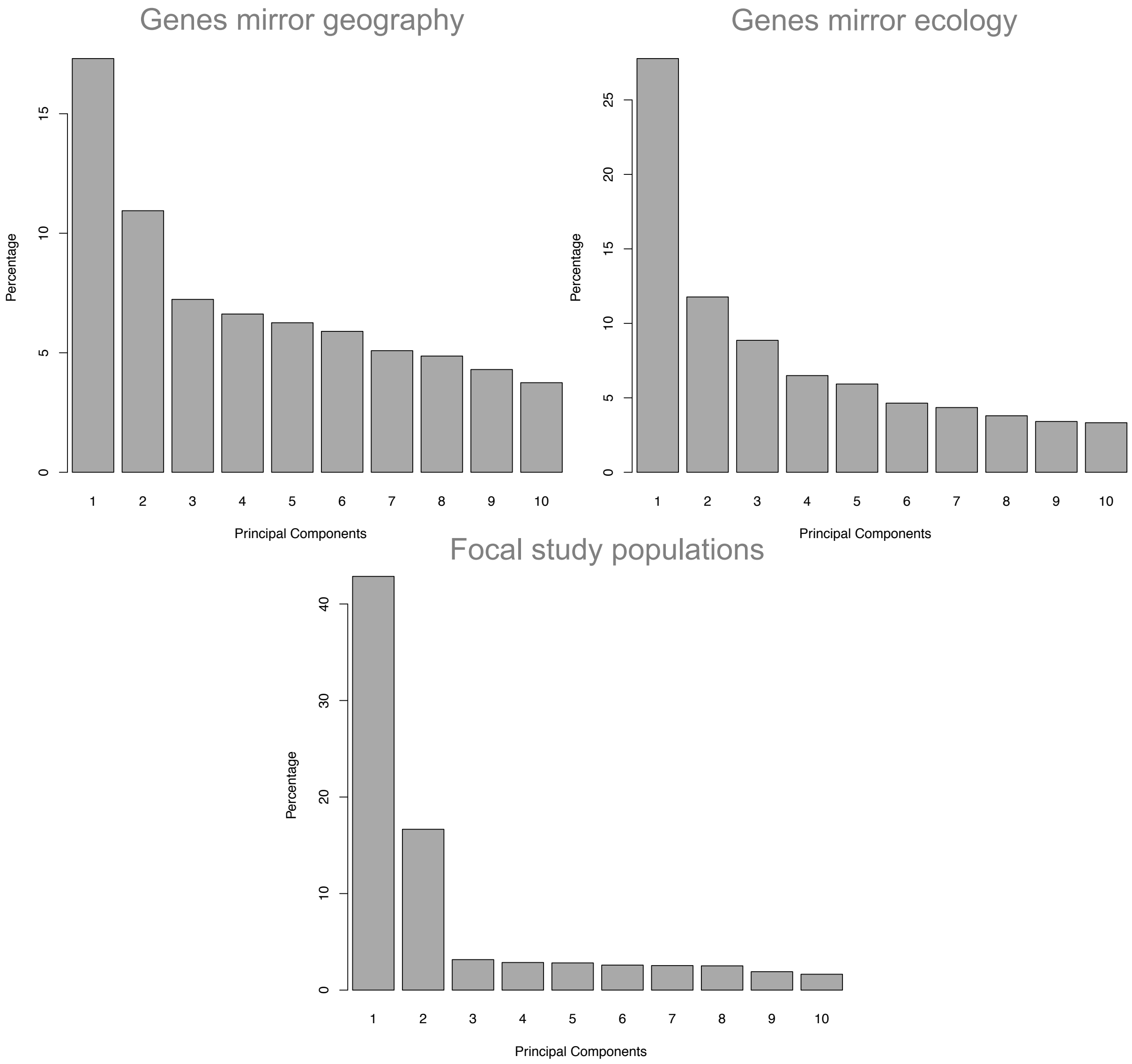


**Figure S3**: Variance of the data explained by eigen vectors 1-10 of PCAs in Figure 2A-C.

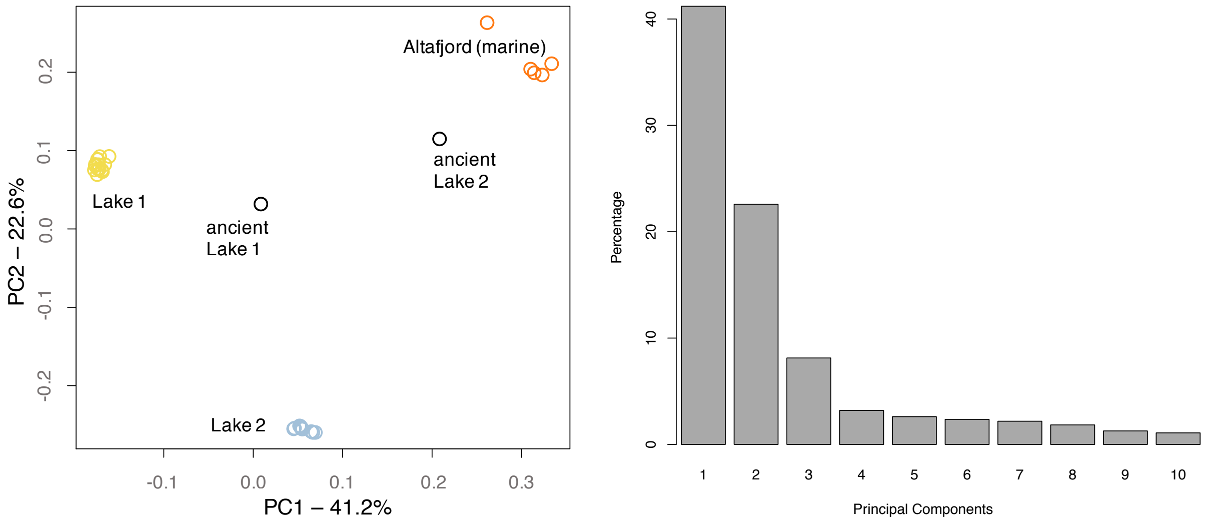


**Figure S4:** (A) Principal component analyses (PCA) of local present-day and ancient samples using transversions in freshwater-marine divergent regions identified by Jones *et al*. (*8*) The first four PCs are significant for $P < 0.05$. (B) Variance of the data explained by eigen vectors 1-10 of PCA in (A).

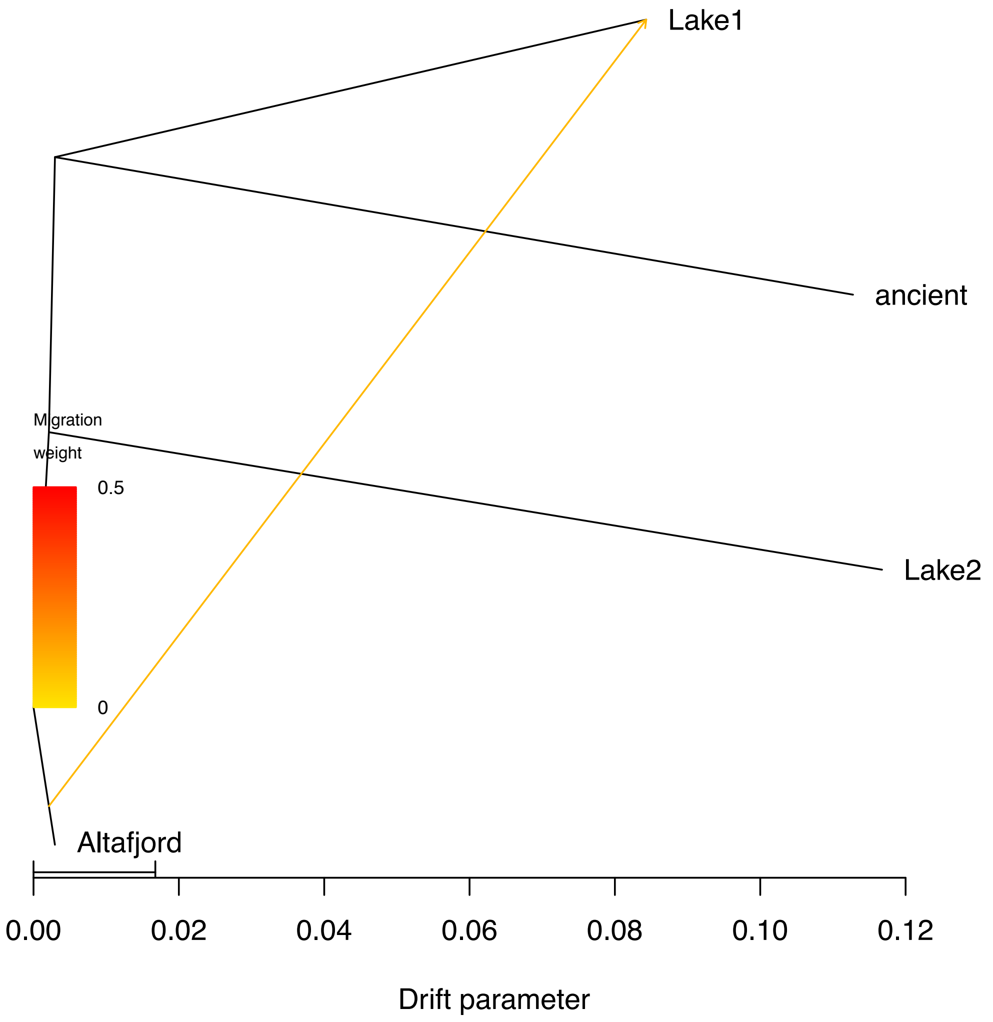


**Figure S5:** TreeMix maximum likelihood graph (*71*) based upon 2,267 transversions. Modern populations are comprised of five high coverage samples. Horizontal branch lengths are proportional to the amount of genetic drift that has occurred along that branch. Note that the long branch of the ancient sample from Lake 2 is an artifact of it being a single pseudo-haploid sample, a pattern typical of low coverage ancient samples (*72*,*73*). However, this does not affect its relative covariance and hence branching order with the modern populations. The scale bar shows 10 times the average s.e. of the entries in the sample covariance matrix. The migration edge inferred using TreeMix is depicted as an arrow colored by migration weight.

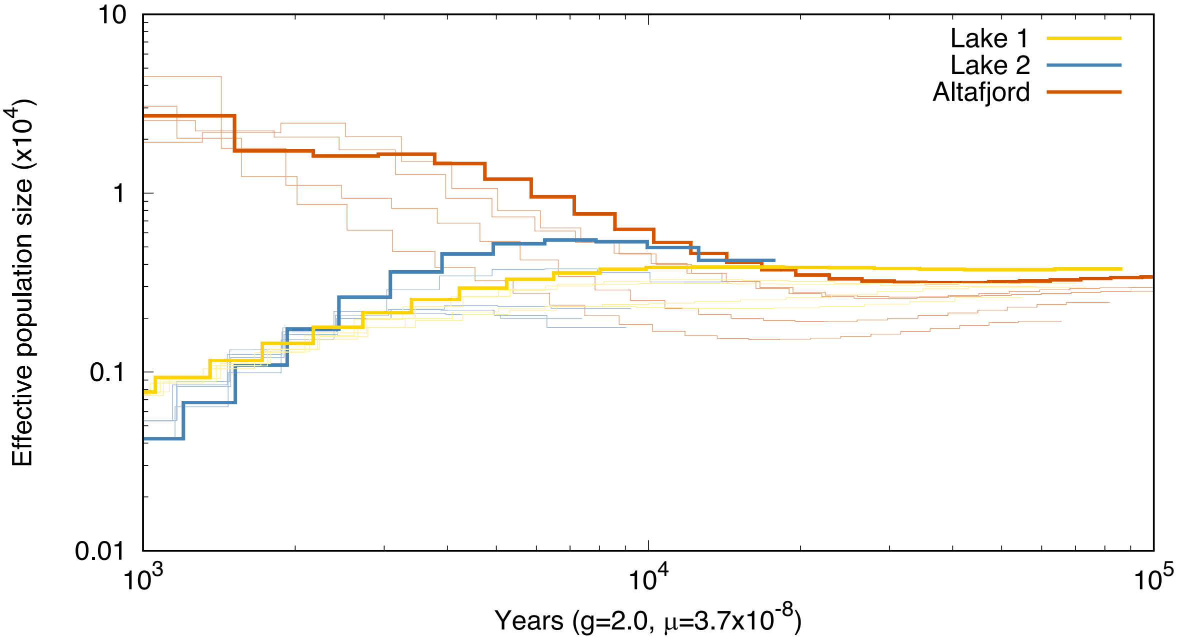


**Figure S6**: PSMC analysis for five highest coverage individuals from each of the three populations. Thick lines represent the genomes included in Fig. 3A.

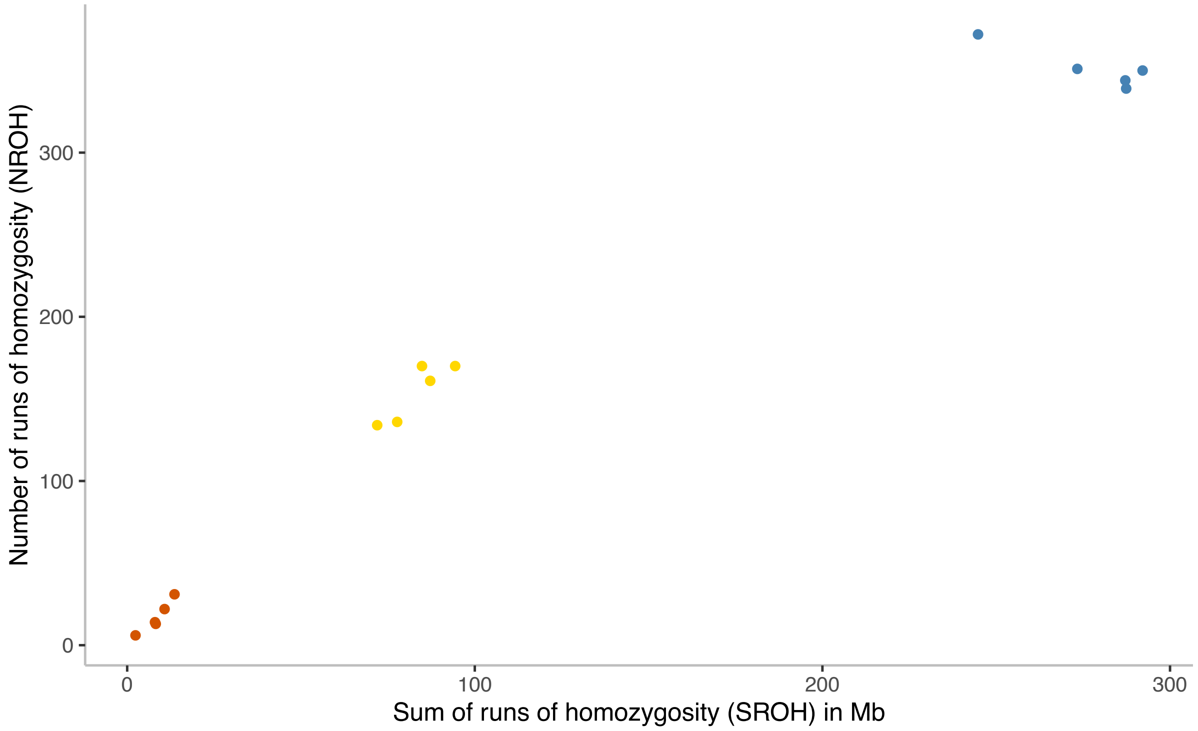


**Figure S7**: ROH analysis for five highest coverage individuals from each of the three populations. Individuals from Lake 1 and Lake 2 are represented in yellow and blue respectively, whereas data of individuals from Altafjord is shown in red.

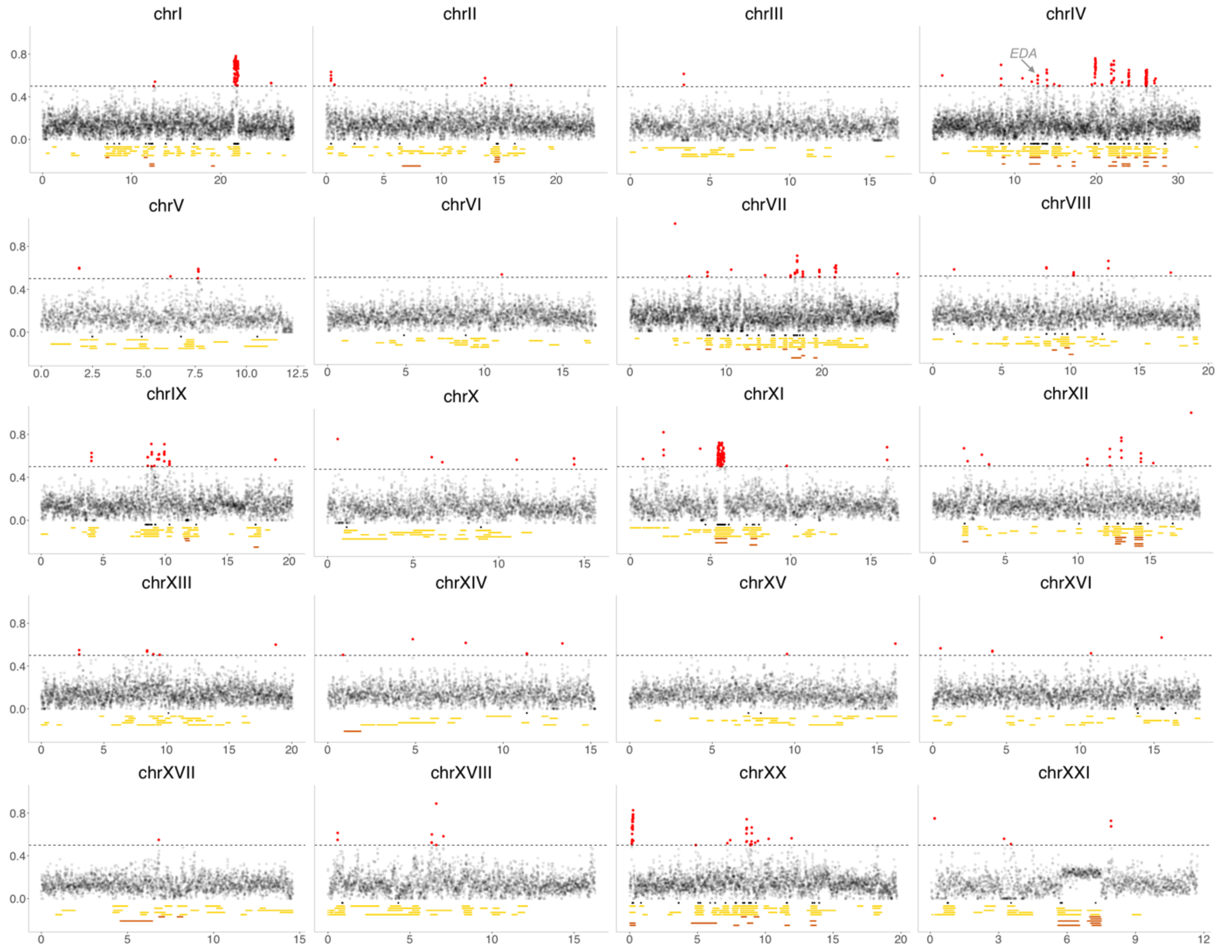


**Figure S8:** Manhattan plots of the genome-wide autosomal $F_{ST}$ analyses between Lake 1 and Altafjord including marked positions of divergent regions and ROH for five individuals of each Lake 1 and Altafjord. The upper part of each plot shows a Manhattan plot for the $F_{ST}$ values across each chromosome with a window size of 10 kb and a sliding window size of 5 kb. The steps on the x-axis represent Mb along the chromosome. Values above 0.5 are coloured red and a dotted marker line is inserted at 0.5. The black bars underneath the Manhattan plot show the location of the 242 divergent regions from Jones et al. (*8*) (FDR 0.05), whereas yellow and red bars represent the locations of the ROHs > 0.3 Mb of five individuals from Lake 1 and Altafjord, respectively.

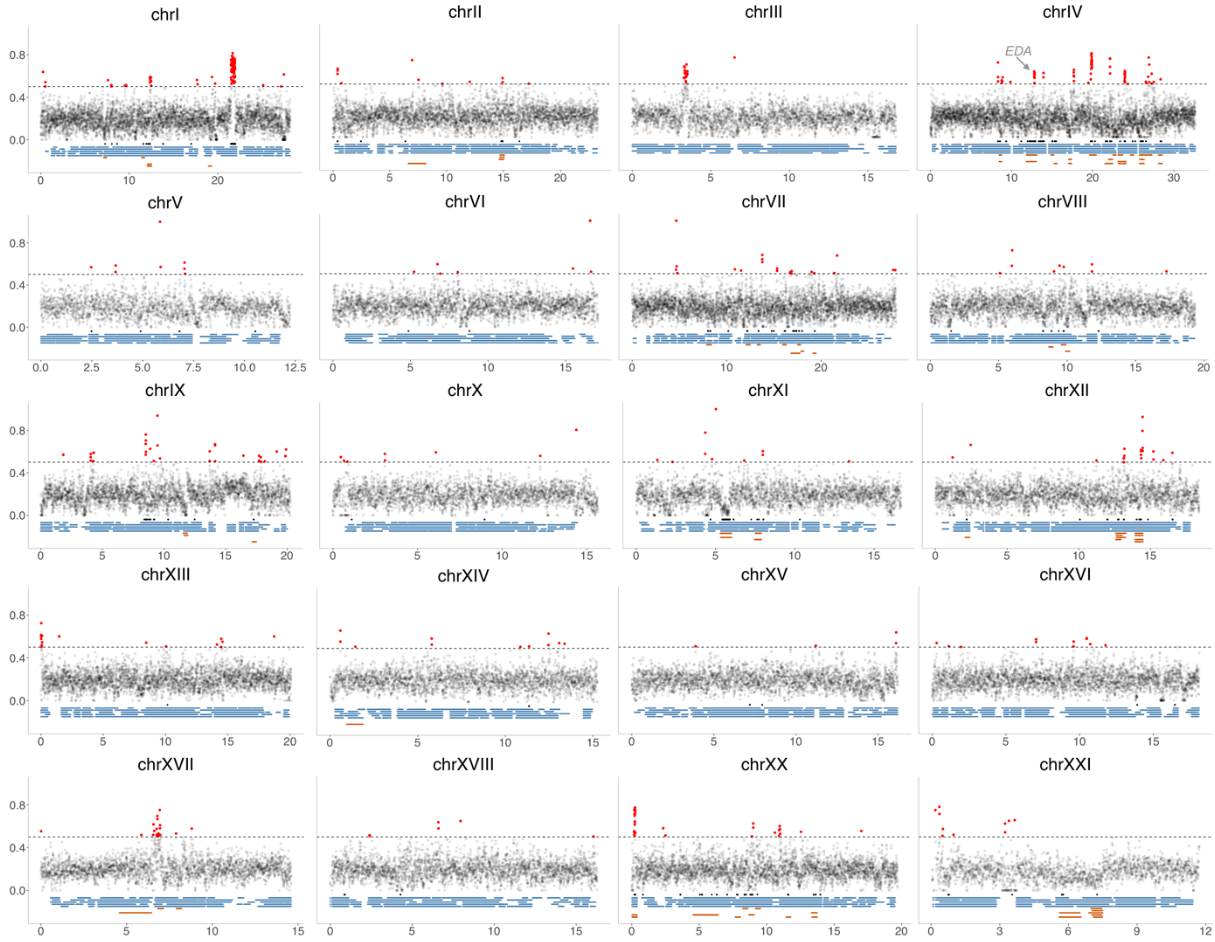


**Figure S9:** Manhattan plots of the genome-wide autosomal $F_{ST}$ analyses between Lake 2 and Altafjord including marked positions of divergent regions and ROH for five individuals of each Lake 2 and Altafjord. The upper part of each plot shows a Manhattan plot for the $F_{ST}$ values across each chromosome with a window size of 10 kb and a sliding window size of 5 kb. The steps on the x-axis represent Mb along the chromosome. Values above 0.5 are coloured red and a dotted marker line is inserted at 0.5. The black bars underneath the Manhattan plot show the location of the 242 divergent regions from Jones et al. (*8*) (FDR 0.05), whereas blue and red bars represent the locations of the ROHs > 0.3 Mb of five individuals from Lake 2 and Altafjord, respectively.

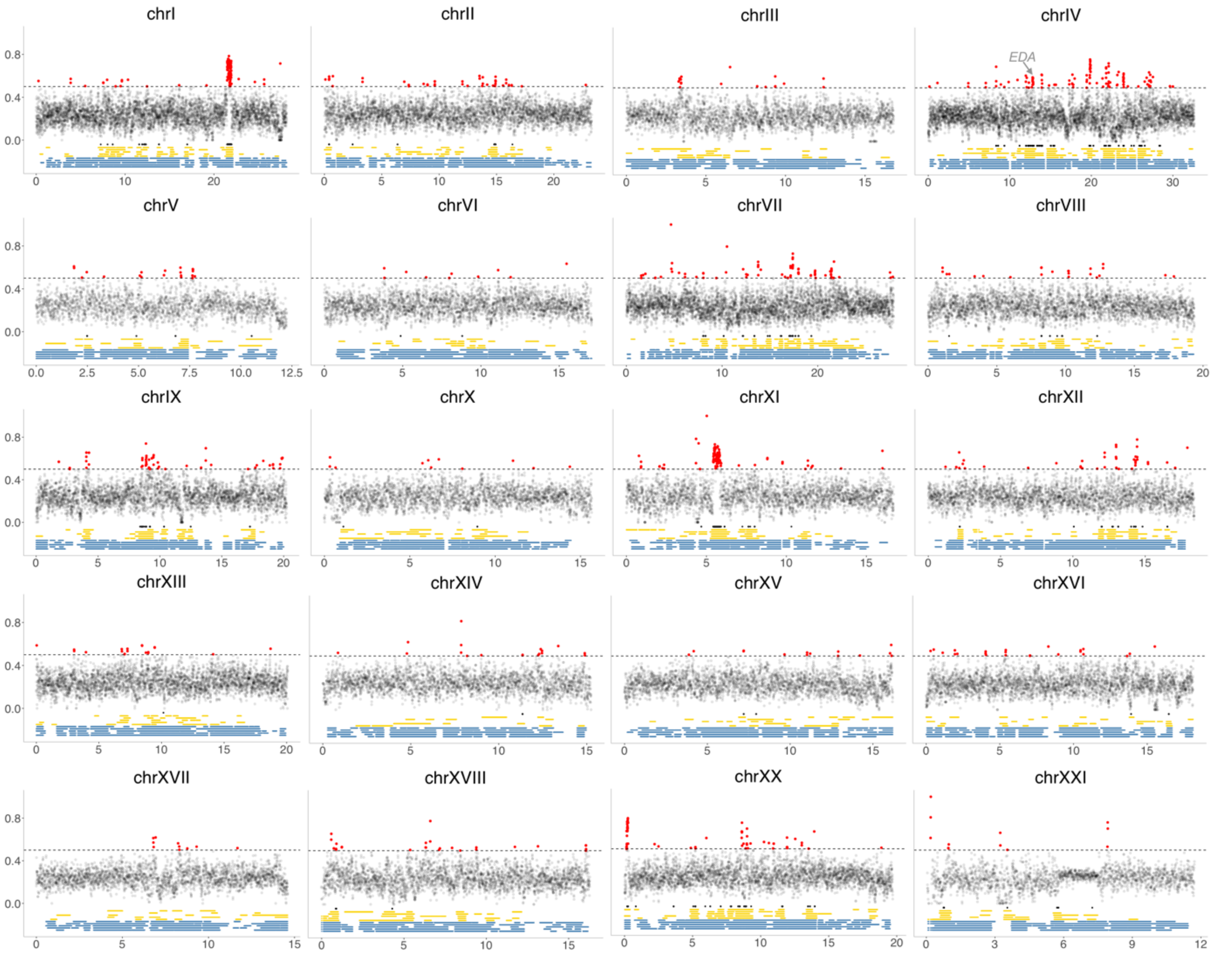


**Figure S10:** Manhattan plots of the genome-wide autosomal $F_{ST}$ analyses between Lake 1 and Lake 2 including marked positions of divergent regions and ROH for five individuals of each Lake 1 and Lake 2. The upper part of each plot shows a Manhattan plot for the $F_{ST}$ values across each chromosome with a window size of 10 kb and a sliding window size of 5 kb. The steps on the x-axis represent Mb along the chromosome. Values above 0.5 are coloured red and a dotted marker line is inserted at 0.5. The black bars underneath the Manhattan plot show the location of the 242 divergent regions from Jones et al. (*8*) (FDR 0.05), whereas yellow and blue bars represent the locations of the ROHs > 0.3 Mb of five individuals from Lake 1 and Lake 2, respectively.

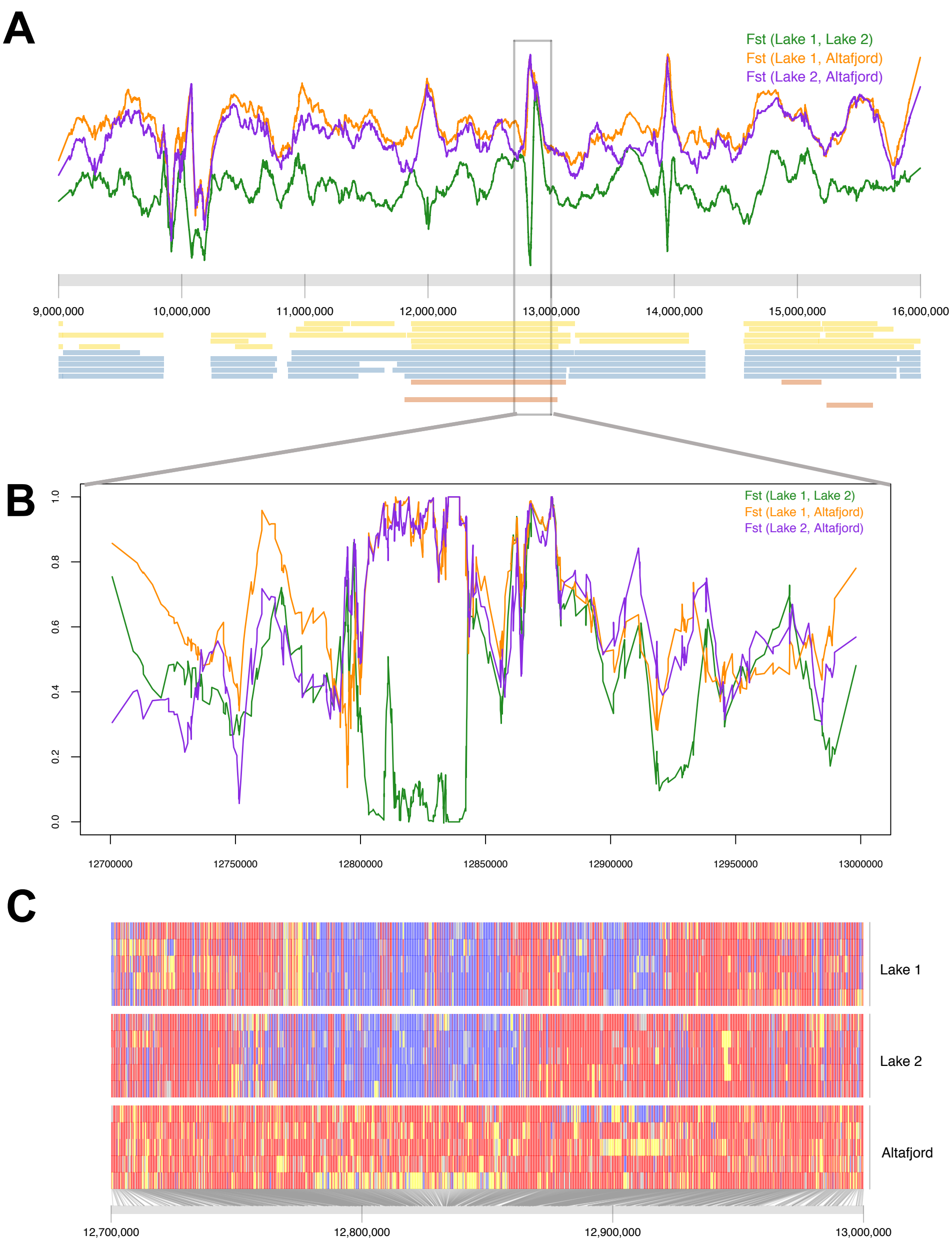


**Figure S11: Freshwater populations carry different EDA haplotype.** (**A**) $F_{ST}$ analyses and ROH for Lake 1, Lake 2 and Altafjord on chrIV:9,000,000-16,000,000 around the EDA region. The upper part shows curves for the $F_{ST}$ values between each two populations, whereas the yellow, blue and red bars underneath the plot represent the locations of the ROHs > 0.3 Mb of five individuals from Lake 1, Lake 2 and Altafjord, respectively. (**B**) $F_{ST}$ analyses for Lake 1, Lake 2 and Altafjord on chrIV:12,700,000-13,000,000 around the focal EDA region. The curves show $F_{ST}$ values between each two populations. (**C**) Underlying genotypes at 1200 randomly picked single nucleotide polymorphisms within chrIV:12,700,000-13,000,000. Rows represent individual fish; columns represent individual single nucleotide polymorphisms; red boxes indicate alleles most common in the marine population; blue boxes indicate alleles less common in the marine population; grey boxes are missing data.

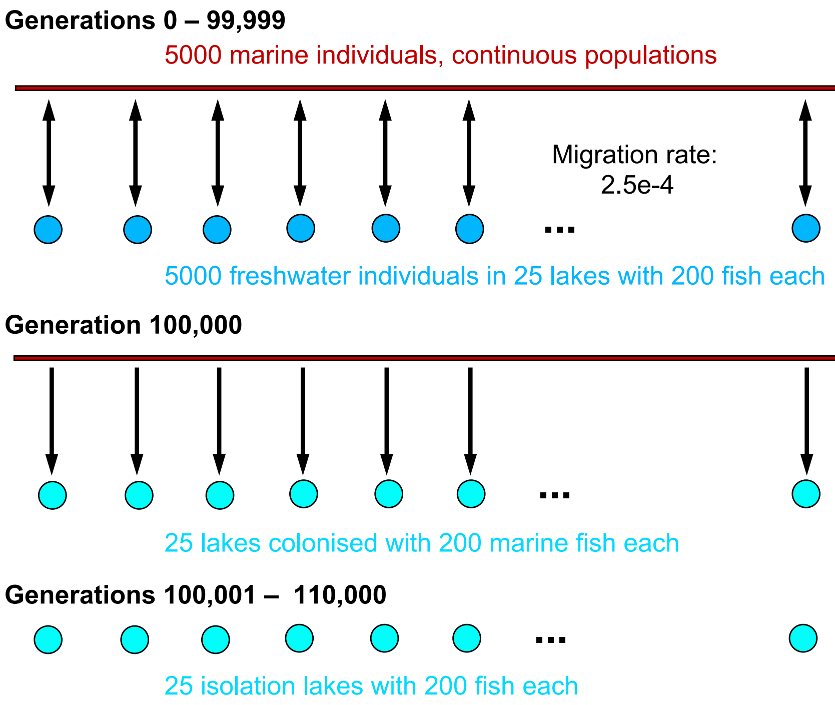


**Figure S12:** Forward **simulations** were performed using SLiM 3 (*74*) based on the elegant model published in Galloway *et al.* (*29*). During the first 100,000 generations two populations, a continuous marine population and a freshwater population separated into 25 lakes, were used to create an initial standing genetic variation of freshwater-adaptive alleles in the marine population. Each population consists of 5,000 individuals and the migration rate between the two populations is $m = 0.00025$. After 100,000 generations, twenty-five new lakes are colonized by 200 marine individuals each. In contrast to the Galloway model, these lakes were modelled as separate populations and were isolated from the marine population, i.e. no migration between the newly introduced lakes and the marine population, or between lakes. The Galloway study simulate a single chromosome of length 1 Gb. In order to study the effects of different recombination rates, we simulate two chromosomes of length 500 Mb with recombination events occurring at a rate of $10^{-8}$ and $10^{-7}$ per bp per generation respectively. The mutation rate in this simulation amounts to $10^{-10}$ per bp per generation. As per the Galloway study, 10 effect regions are spread across the chromosome(s). Within these effect regions, mutations of six different types can arise. Mutations can either be dominant (dom), recessive (rec) or additive (add). Furthermore, each mutation is either beneficial in the marine environment and deleterious in the freshwater environment (mar), or vice versa (fw). The effect sizes for each mutation are chosen randomly from an exponential distribution with a mean ±0.5. Individual trait values are subsequently determined additively from the diploid genotypes and individual fitness is calculated from these individual trait values as in Galloway *et al.* (*29*). SLiM code is available at https://github.com/Stickle-Back-in-Time/Stickle-Back-in-Time.

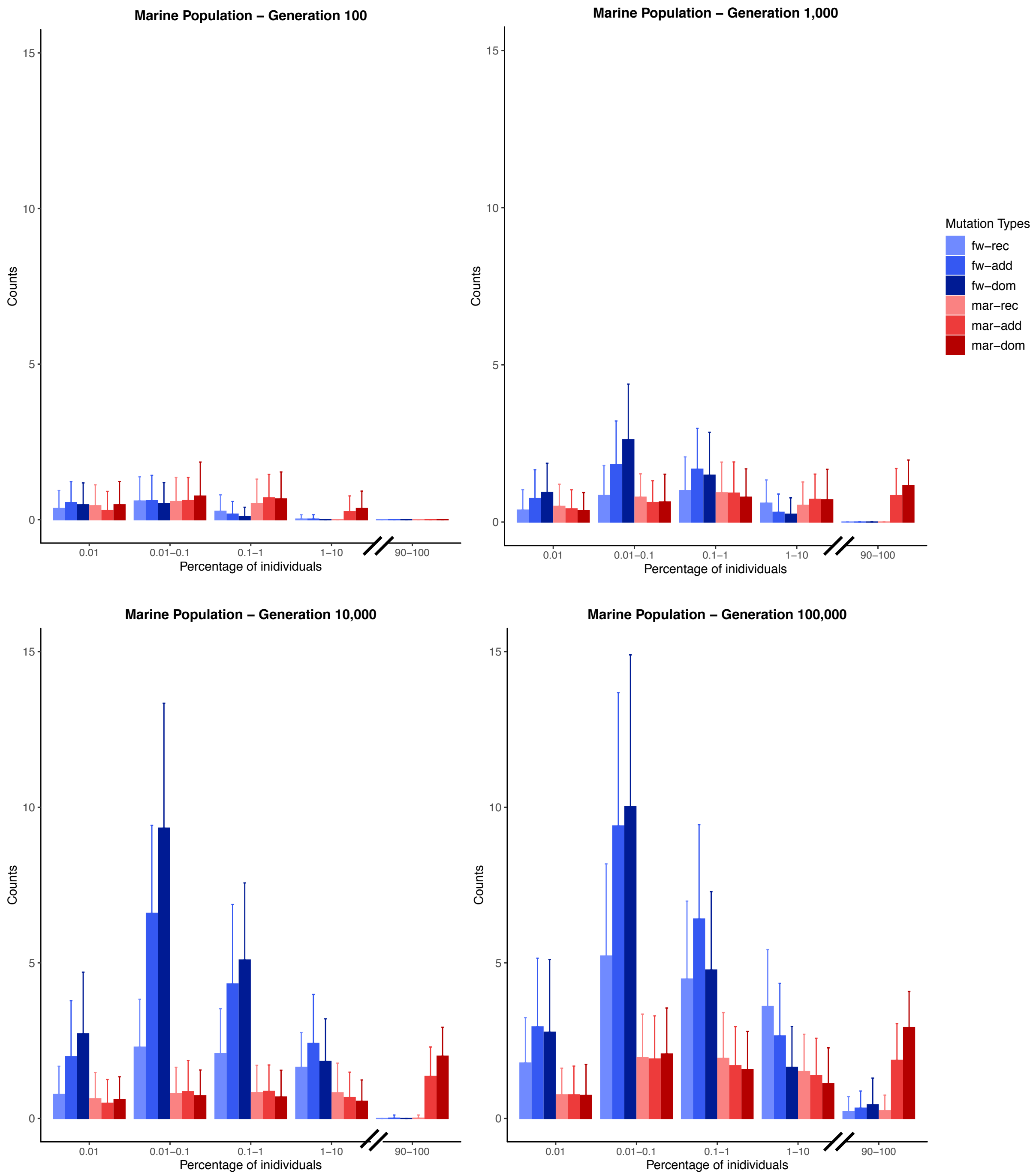


**Figure S13:** Counts of different mutation types in the marine population at generation 100, 1,000, 10,000 and 100,000 – means and standard deviation from 100 simulation runs. Dominant and additive mutations that are beneficial in the marine environment (red) rise to high frequencies over time, mutations that are beneficial in freshwater but deleterious in the marine environment (blue) stay at low frequencies (mostly < 0.1, which corresponds to 10 haplotypes in the whole marine population) and represent standing genetic variation of freshwater-adaptive mutations.

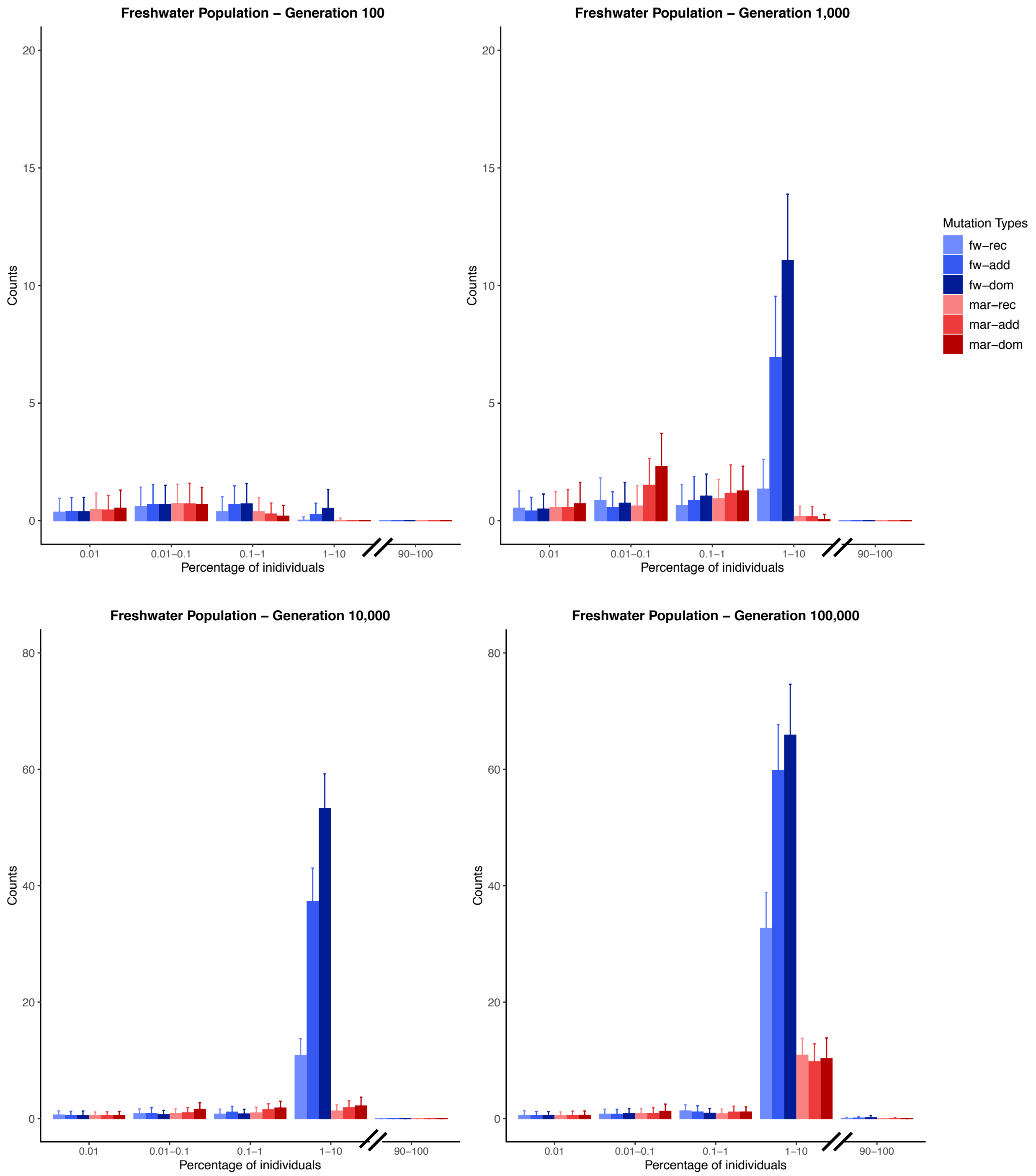


**Figure S14:** Counts of different mutation types summed up for all freshwater lakes at generation 100, 1,000, 10,000 and 100,000 – means and standard deviation from 100 simulation runs. Most mutation types occur at a frequency less than 10%, which corresponds to 1,000 haplotypes of all 10,000 haplotypes of freshwater individuals. A mutation at a frequency of 10% in the freshwater population could be fixed in two freshwater lakes.

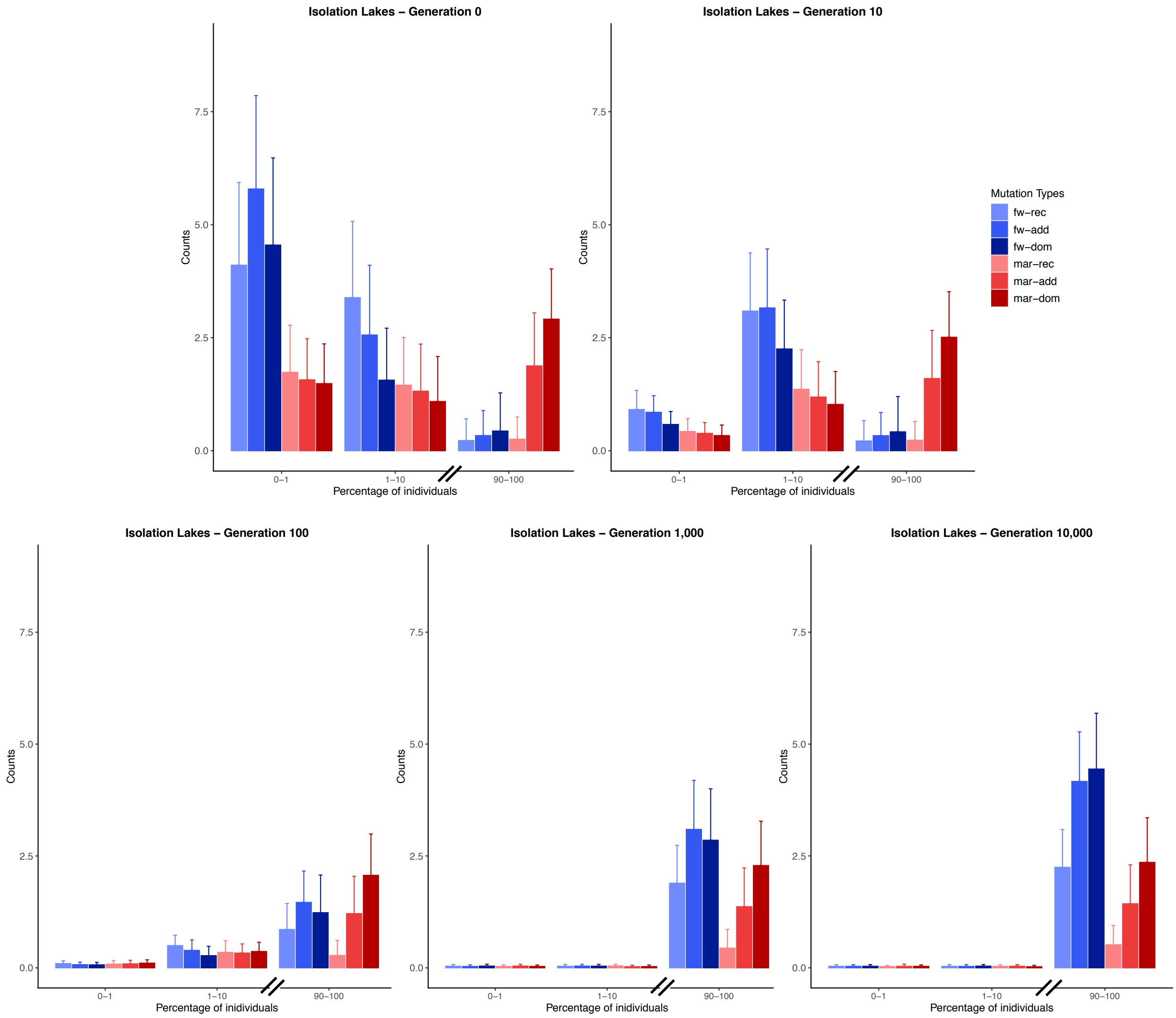


**Figure S15:** Counts for different mutation types for all isolation lakes at 0, 10, 100, 1,000 and 10,000 generations after colonization from the marine environment – means and standard deviation from 100 simulation runs. Marine-adaptive alleles (red) introduced into the isolation lakes at high frequencies during colonization are retained as fixed substitutions. Freshwater-adaptive mutations (blue) rise to high frequency over time.

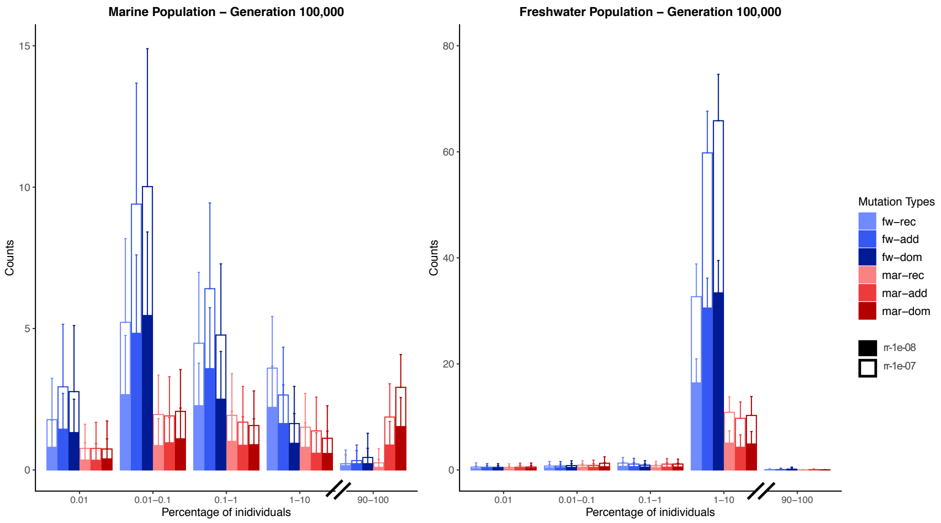


**Figure S16:** Stacked counts for different mutation types showing the effect of different recombination rates for marine and freshwater population at generation 100,000 – means and standard deviation from 100 simulation runs. Mutations rising to high frequency appear at similar abundance on the two chromosomes with one chromosome having a recombination rate of 1e-08 (rr-1e-08) and the other 1e-07 (rr-1e-07).

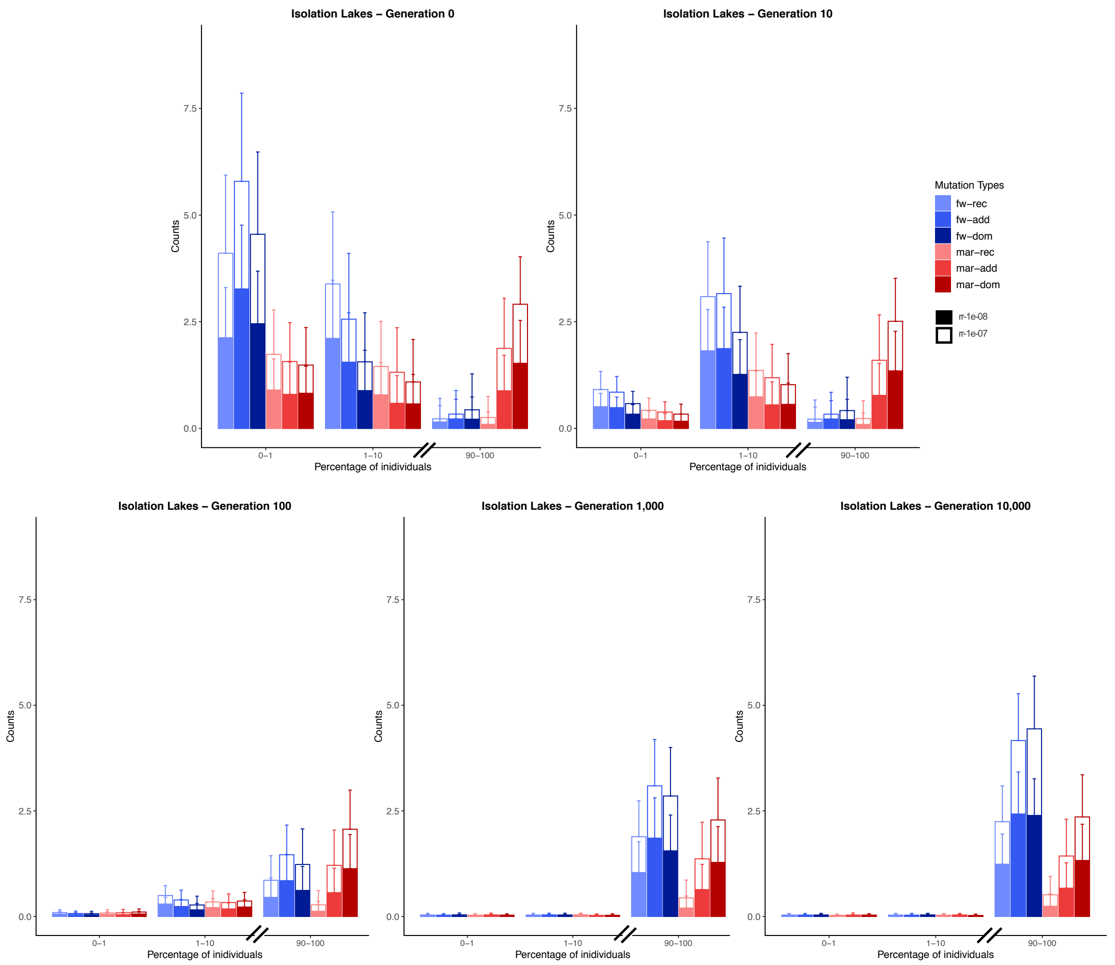


**Figure S17:** Stacked counts for different mutation types and different recombination rates for all isolation lakes at generation 0, 10, 100, 1,000 and 10,000 after colonization at generation 100,000 – means and standard deviation from 100 simulation runs. Mutations rising to high frequency appear at similar abundance on the two chromosomes with one chromosome having a recombination rate of 1e-08 (rr-1e-08) and the other 1e-07 (rr-1e-07).

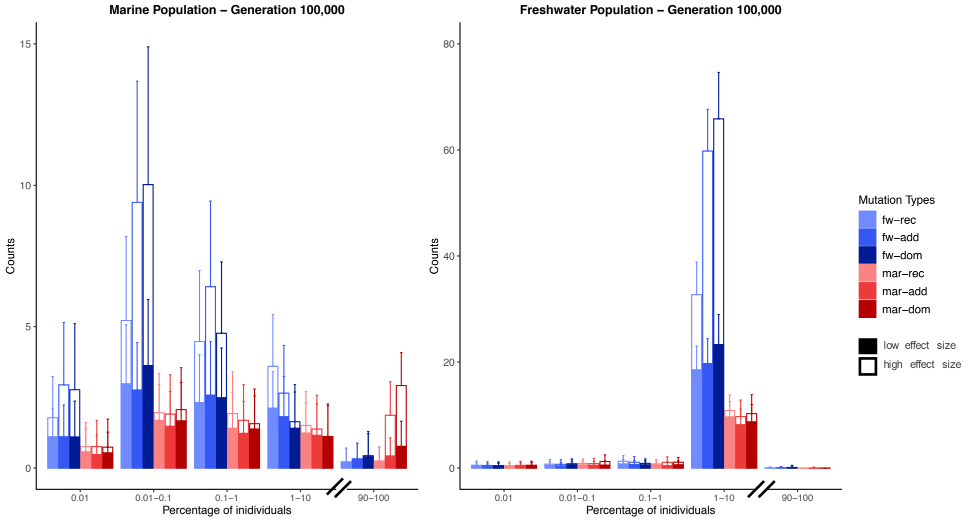


**Figure S18:** Stacked counts for different mutation types showing the effect of different effect sizes for marine and freshwater population at generation 100,000 – means and standard deviation from 100 simulation runs. Dominant and additive mutations that are beneficial in the marine environment (red) and at high frequencies (90-100%) in the marine population have mostly a large effect size (> 0.5), whereas these mutation classes with small effect sizes (< 0.5) are more common at low frequencies. Respectively, mutations beneficial in a freshwater environment (blue) at intermediate frequency (here 1-10%) in the freshwater population show large effect sizes (> 0.5), whereas those beneficial in marine environments (red) have mostly low effect sizes (< 0.5).

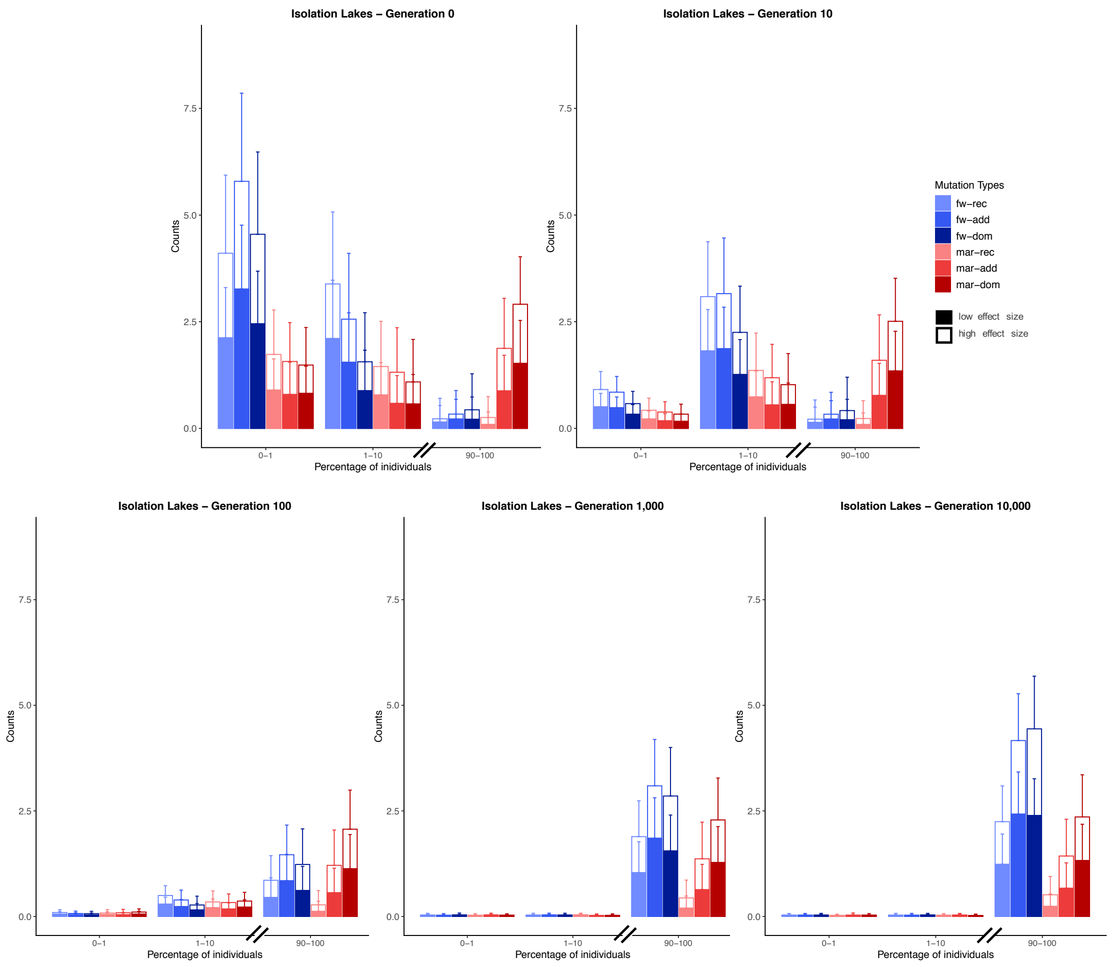


**Figure S19:** Stacked counts for different mutation types and different effect sizes for all isolation lakes at generation 0, 10, 100, 1,000 and 10,000 after colonization at generation 100,000 – means and standard deviation from 100 simulation runs. Whereas marine mutations (red) decrease in frequency over time, all types of freshwater mutations (blue) rise to high frequency. These mutations show high (> 0.5) and low effect sizes (< 0.5) in a similar abundance.

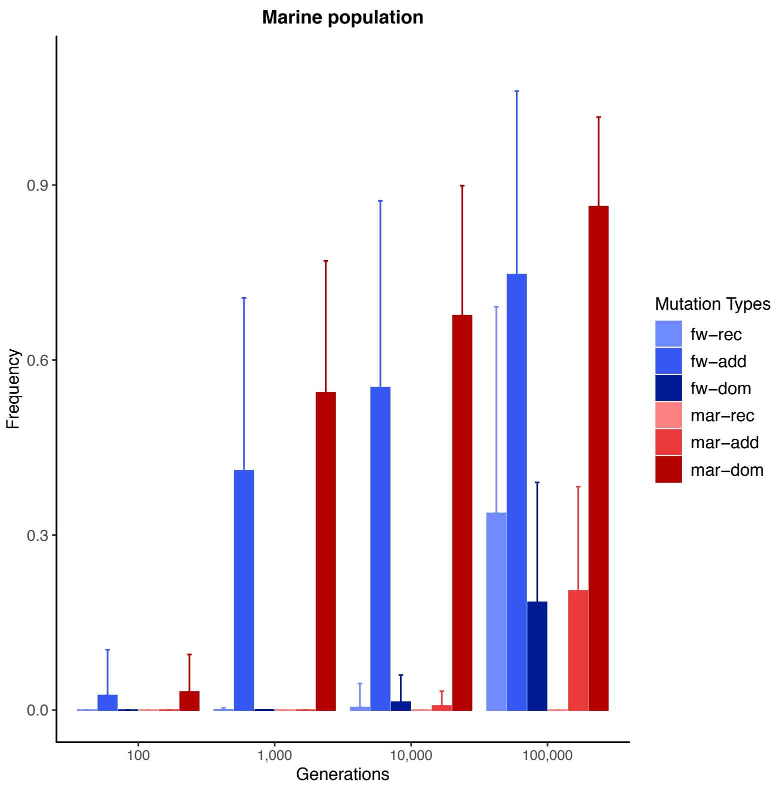


**Figure S20:** Mean frequency of all mutations from each mutation type in the marine population at generation 100, 1,000, 10,000 and 100,000 for mutations being present in at least one of the isolation lakes until generation 110,000 – means and standard deviation from 100 simulation runs. Marine-dominant as well as freshwater-additive mutations are at high frequencies, whereas freshwater-recessive, freshwater-dominant, marine-recessive and marine-additive mutations are at low frequencies at generation 100,000 in the marine population.

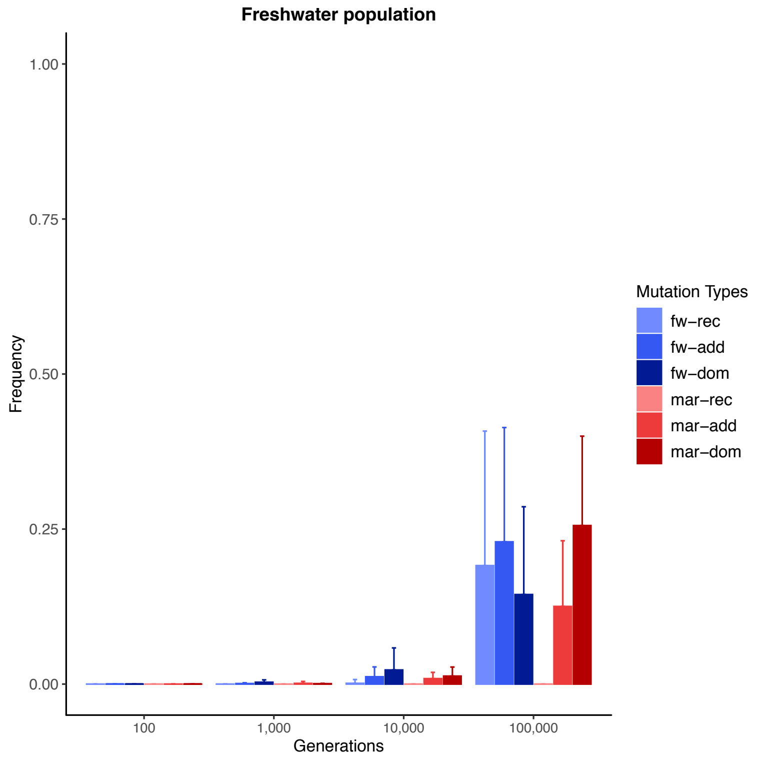


**Figure S21:** Mean frequency of all mutations from each mutation type in the freshwater population at generation 100, 1,000, 10,000 and 100,000 for mutations being present in at least one of the isolation lakes until generation 110,000 – means and standard deviation from 100 simulation runs. Freshwater as well as marine mutations are present at frequencies between 0 and 0.4 in the freshwater population.

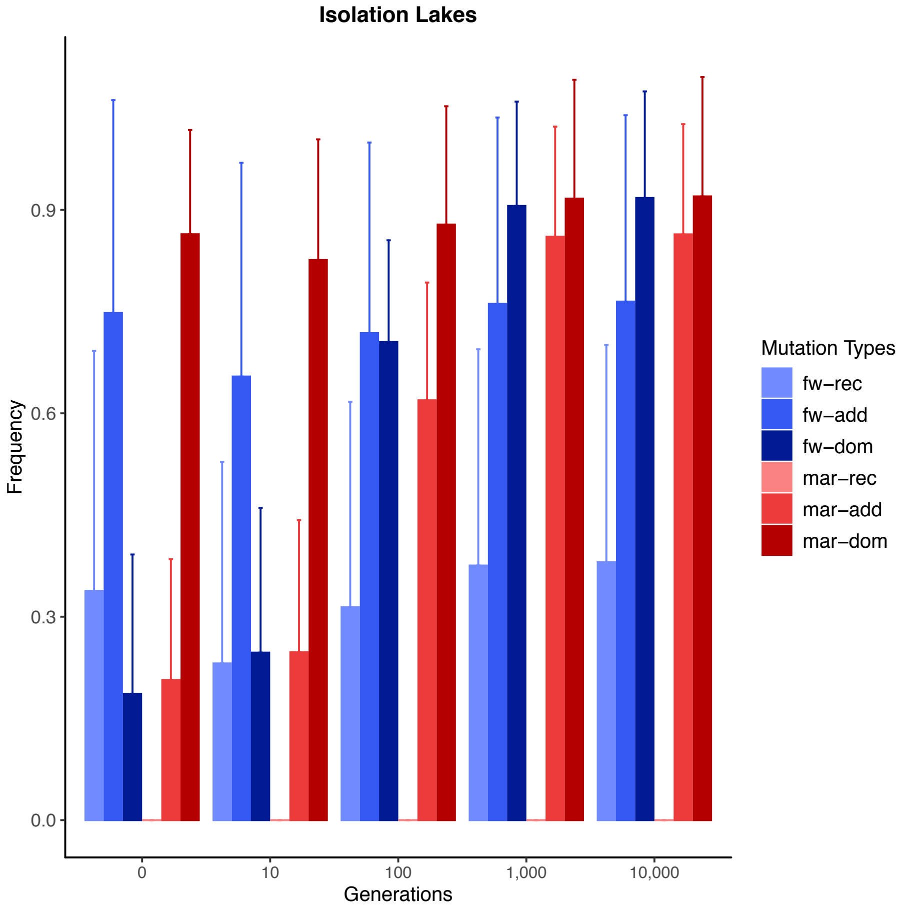


**Figure S22:** Mean frequency of all mutations from each mutation type in the isolation lakes at generation 0, 10, 100, 1,000 and 10,000 for mutations being present in at least one of the isolation lakes until generation 10,000 after colonization at generation 100,000 – means and standard deviation from 100 simulation runs. Frequencies of mutation types at generation 0 are similar to frequencies in the marine population at generation 100,000. The frequency of all mutation types are increasing over time. Whereas, marine dominant alleles start at high frequency at generation 0, freshwater-dominant and marine-additive mutations rise from low to high frequencies over time.

**A**

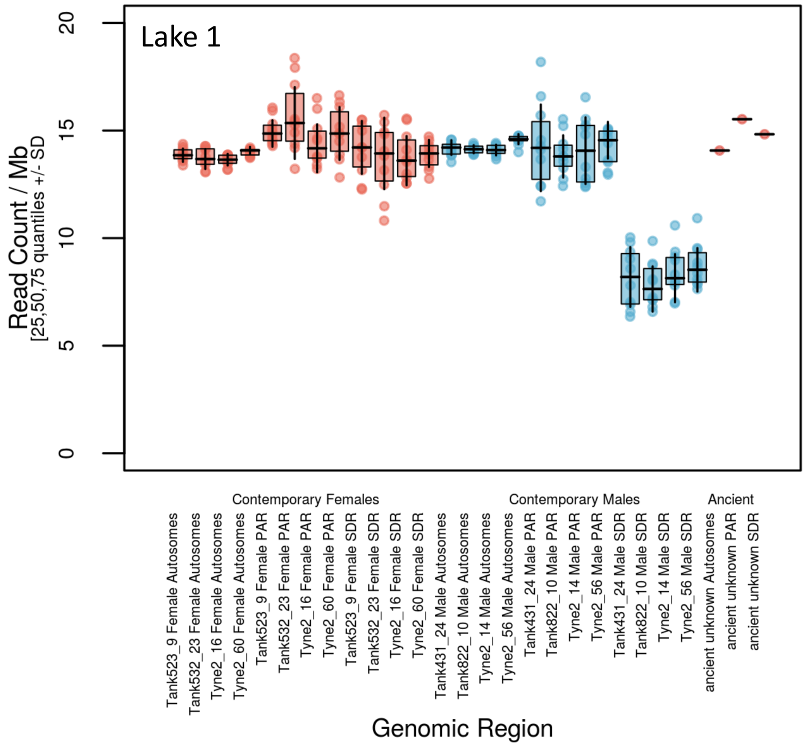


**B**

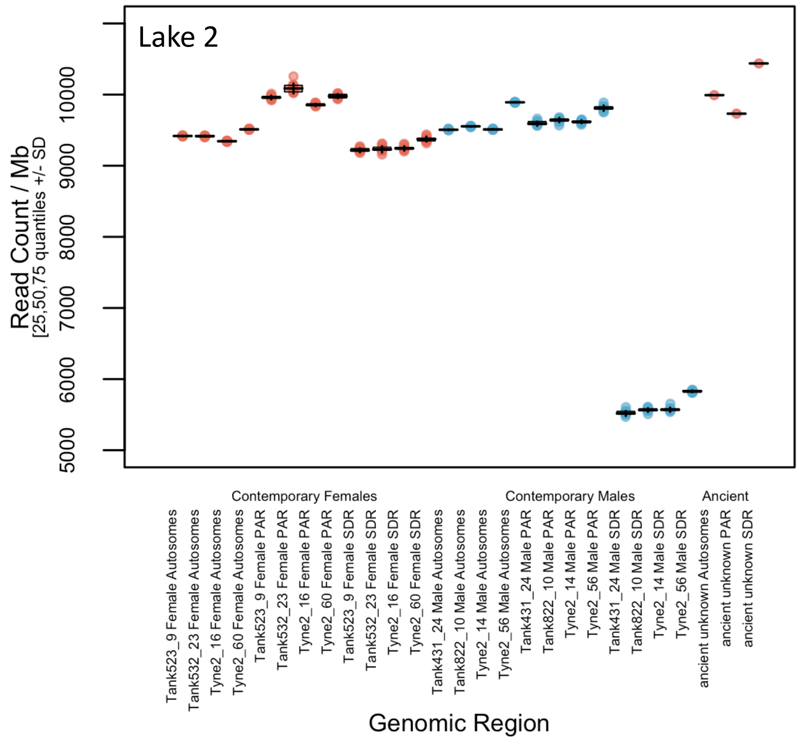


**Figure S23:** Sexing by coverage of **a.** Lake 1 and **b**. Lake 2. Comparison of the mean coverage of the autosomes, the pseudoautosomal region, and the sex-determining region among 4 female (red) and 4 male (blue) modern samples down-sampled to comparable coverage with the ancient sample.

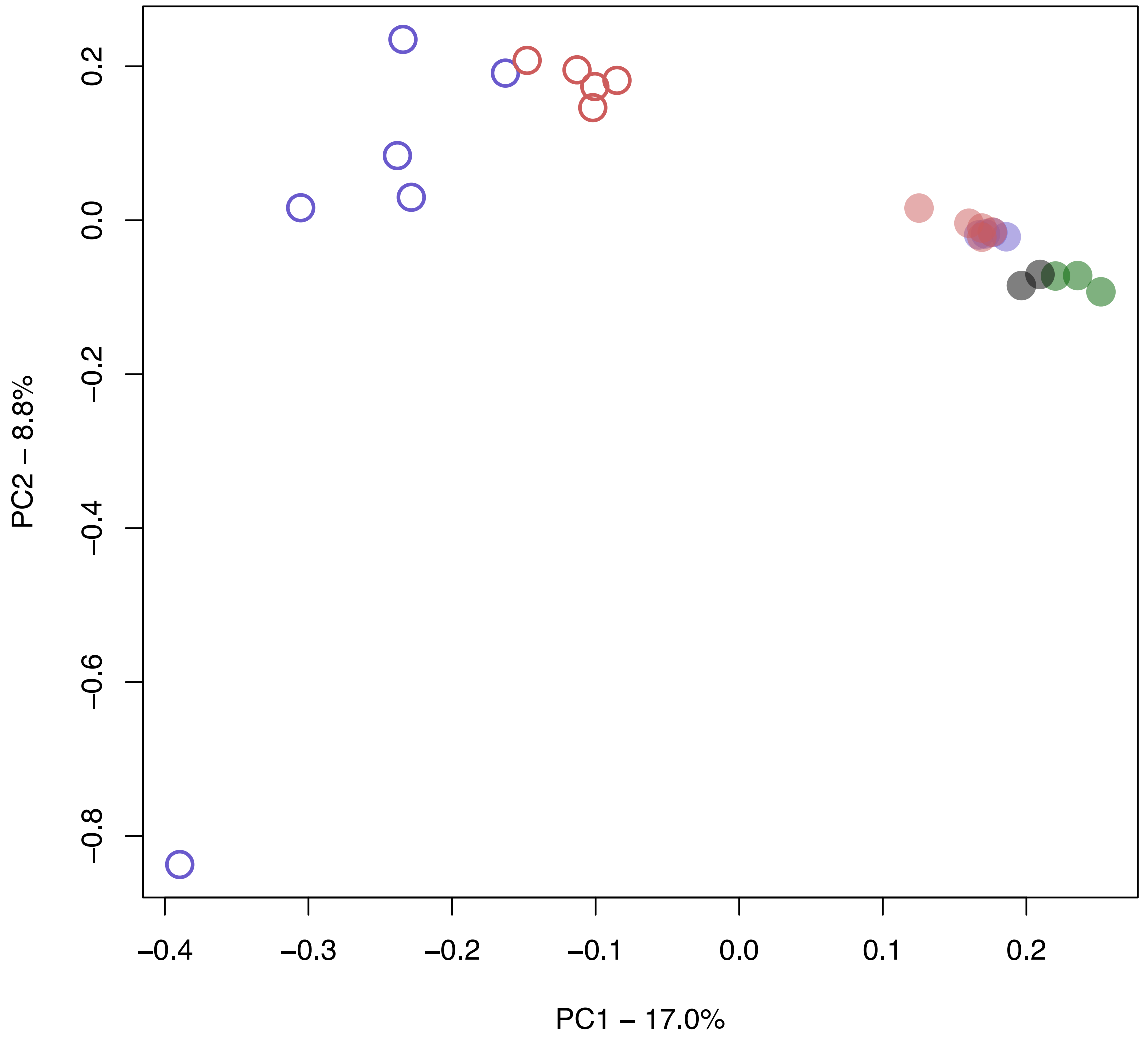


**Figure S24:** Principal component analyses (PCA) of the global dataset from Jones et al. (*8*), one randomly sampled individual each from Altafjord, Lake 1 and Lake 2 (all coloured green) and the ancient samples, based on transversions in non-divergent regions.

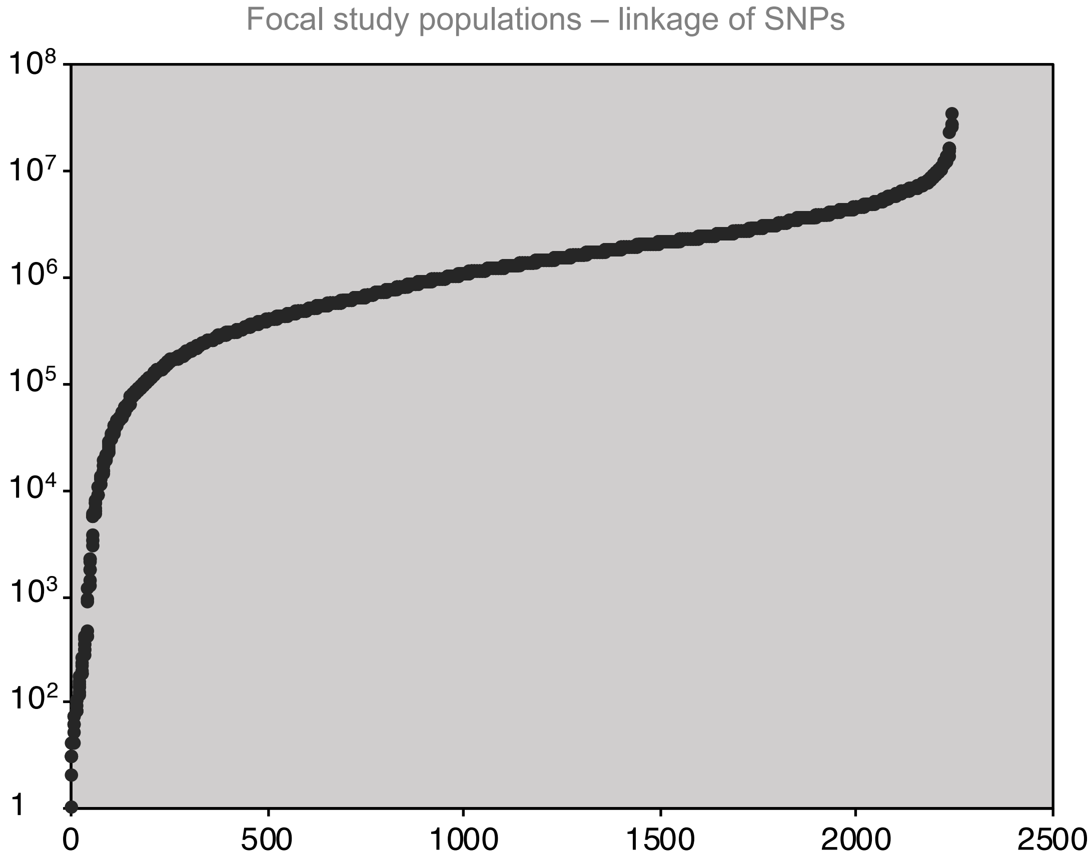


**Figure S25:** SNPs included in PCA comparing Lake 1, Lake 2, Altafjord and the ancient samples in Fig. 2C. The x-axis is the cumulative number of SNPs, ordered based on their spacing with the next nearest SNP in the dataset. The *y*-axis is the distance in base pairs to the next nearest SNP on the same chromosome.

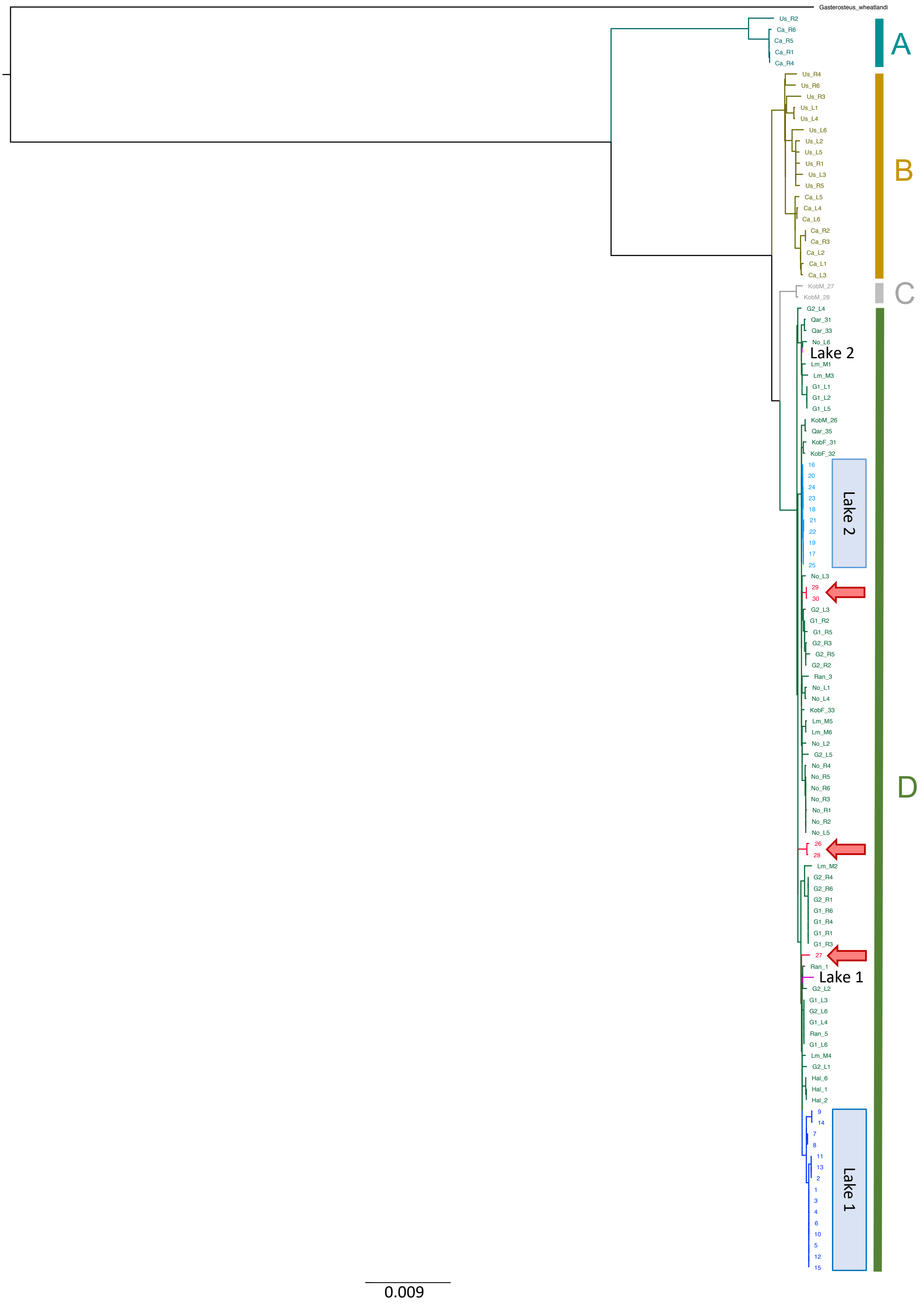


**Figure S26**: Maximum likelihood phylogeny of mitochondrial genomes from samples sequenced by this study combined with the dataset used by Liu et al. (*22*). As per Liu et al., lineage A corresponds to the 'Japanese Lineage', lineage B to the Pacific part of a major 'Euro-American' lineage, lineage C to the 'Transatlantic Lineage' and lineage D to the 'European Lineage'. Modern samples from Lakes 1 and 2 are demarcated by blue boxes, samples from Altafjord by red arrows, and the two ancient samples by the black labels 'Lake 1' and 'Lake 2'.

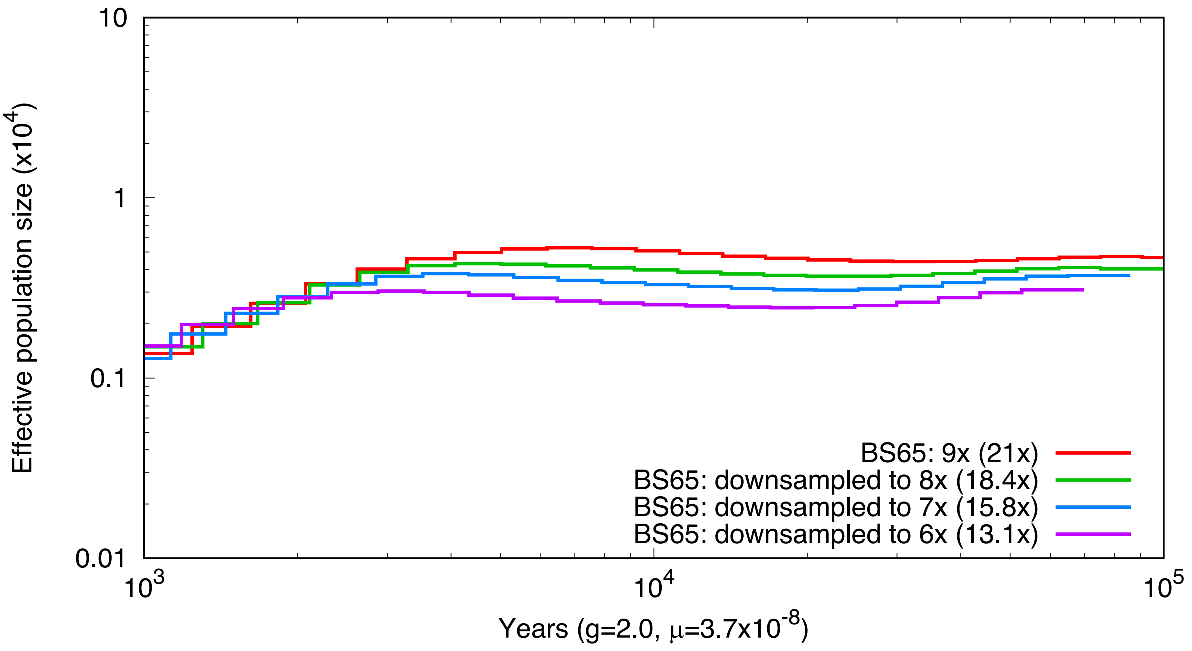


**Figure S27:** PSMC plots for sequencing data from freshwater stickleback BS65 (*60*) downsampled to different coverages.
